# Translational control of innate barrier defense by the gut microbiota

**DOI:** 10.1101/2025.08.11.668728

**Authors:** Yun Li, Runrun Wu, Xianda Ma, Jianglin Zhang, Junjie Ma, Yiran Duan, Brian Hassell, Kelly A. Ruhn, Cassie L. Behrendt, Chaitanya Dende, Li Wang, Andrew Lemoff, Prithvi Raj, Zheng Kuang, Lora V. Hooper

## Abstract

The intestinal epithelium is protected by a mucus barrier, infused with antimicrobial proteins, that restricts microbial access to host tissue. Because assembling this barrier is energetically costly, its production must be tightly regulated. Here, we show that the microbiota regulates the translation of key mucus barrier components, including the structural glycoprotein mucin 2 and the antimicrobial enzyme lysozyme. This process is initiated by microbial induction of histone deacetylase 5 (HDAC5) in secretory epithelial cells of the intestine. HDAC5 promotes deacetylation of 14-3-3 proteins, enabling activation of the energy sensing kinase mTOR which enhances translation of mucus and antimicrobial proteins. These findings reveal a mechanism by which the microbiota controls barrier immunity at the level of protein synthesis and suggest that the HDAC5–mTOR axis integrates microbial and energetic signals to regulate intestinal defense.

**One sentence summary:** The gut microbiota enhances translation of innate barrier defense proteins through HDAC5-mediated activation of mTORC1 signaling.

## Introduction

The intestinal epithelium is a vast body surface that is exposed to the external environment. The large epithelial interface optimizes nutrient absorption but also exposes the host to trillions of microorganisms inhabiting the gut lumen. Intestinal epithelial cells (IECs) secrete a diverse group of innate defense proteins that defend the gut barrier against microbial invasion. These include mucin glycoproteins that assemble into a protective mucus barrier (*1, 2*), and mucus-associated antimicrobial proteins that reinforce this barrier by killing, aggregating, or blocking the attachment of bacteria (*3*). Disrupted production of mucus barrier components can lead to infection, inflammation, and metabolic disease, underscoring the essential role of this barrier in maintaining health (*1, 4, 5*).

The microbiota exerts transcriptional control over the expression of many intestinal barrier defense proteins. For instance, gut microbes induce the transcription of genes encoding the antimicrobial proteins REG3G, RELMβ, and SPRR2A (*6-8*). This tight transcriptional control likely reflects the high energetic cost of producing these proteins, ensuring they are expressed only when necessitated by microbial threats. However, genes encoding a subset of barrier defense proteins, including the mucus glycoprotein mucin 2 and the antimicrobial enzyme lysozyme, can be expressed even in the absence of a microbiota (*9*). This raises the question of how expression of these proteins is coordinated with both physiological demand and energetic constraints.

Here, we show that the gut microbiota regulates the translation of key group of proteins that defend the intestinal surface from microbial invasion. We identify a mechanism that is initiated by microbiota-induced expression of histone deacetylase 5 (HDAC5) in gut secretory epithelial cells. HDAC5 deacetylates the regulatory protein 14-3-3, enabling it to sequester Raptor, a component of the energy-sensing mTORC1 complex. This frees mTOR to activate downstream signaling pathways that promote translation of barrier defense proteins. Our findings thus reveal a mechanism by which the microbiota enhances gut innate immunity through translational control. Because HDAC5 responds to microbial cues and mTOR to cellular energy status, these results suggest that the HDAC5—mTOR axis may integrate microbial and metabolic signals to regulate immune defense in the intestine.

## Results

### HDAC5 promotes expression of barrier defense proteins in intestinal secretory epithelial cells

Histone deacetylases (HDACs) have emerged as key regulators of host–microbiota interactions. For example, HDAC3 promotes microbiota-dependent homeostasis, lipid metabolism, and epithelial repair. It is regulated by the microbiota both transcriptionally and through modulation of its enzymatic activity (*10-13*). Given that the gene encoding HDAC5 is also highly expressed in intestinal epithelial cells (*13*), we asked whether its expression is similarly influenced by the microbiota. *Hdac5* transcript levels were significantly higher in conventional compared to germ-free mice (Fig. 1, A and B, and fig. S1A). This microbiota-dependent increase was also evident at the protein level (Fig. 1C; fig. S1, B and C), indicating that the gut microbiota promotes HDAC5 expression in the intestinal epithelium.

**Figure 1.**
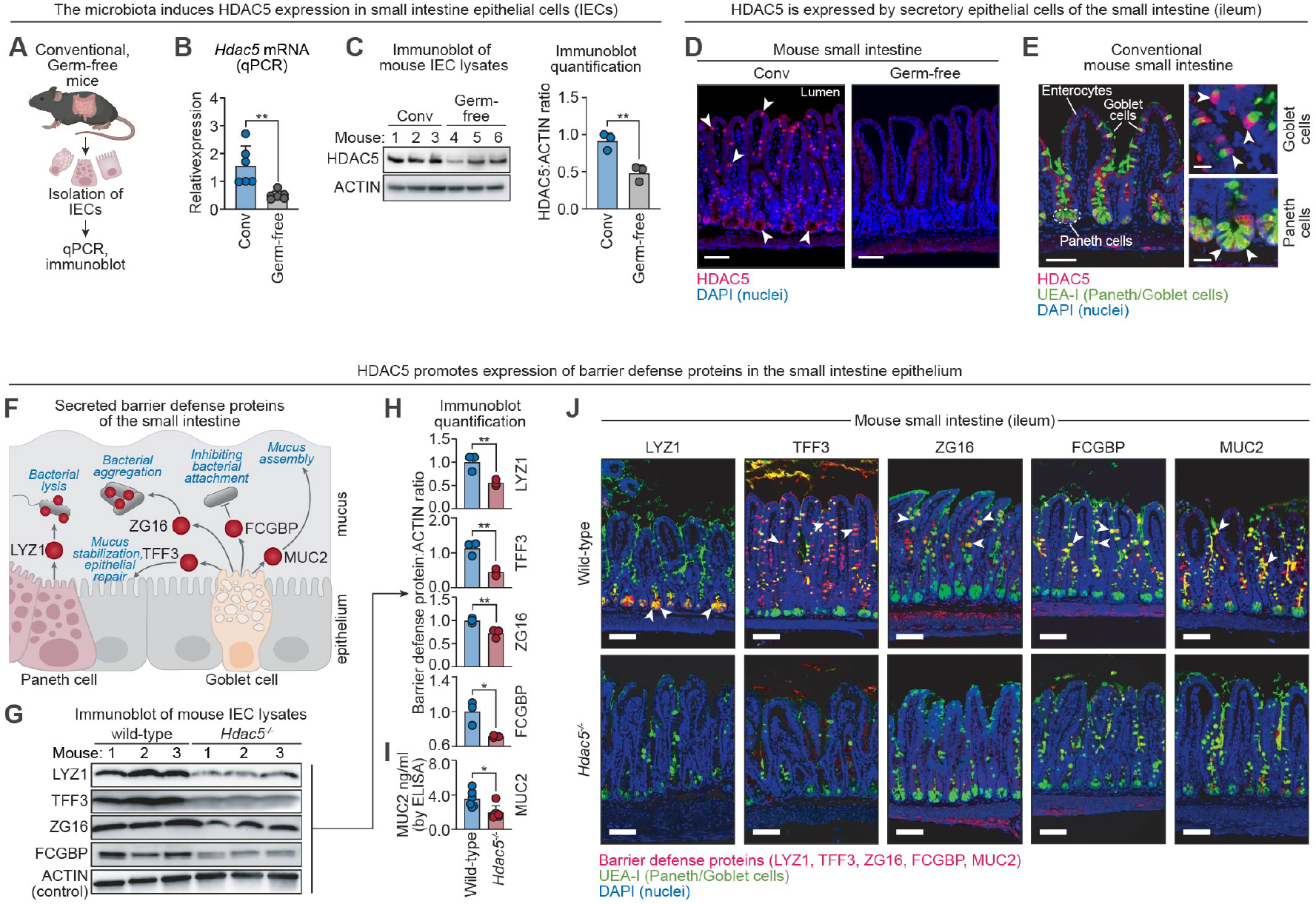
HDAC5 promotes expression of barrier defense proteins in intestinal secretory epithelial cells. **(A)** Experimental overview: epithelial cells were isolated from the small intestines of conventional (conv) and germ-free mice by EDTA treatment and were analyzed by real-time quantitative PCR (qPCR) and immunoblot. **(B)** qPCR analysis of *Hdac5* transcripts in small intestinal epithelial cells of conventional and germ-free mice. **(C)** Immunoblot detection of HDAC5 in small intestine epithelial cell (IEC) lysates from conventional and germ-free mice. ACTIN was the loading control. Protein band intensities were measured by scanning densitometry and ratios were calculated. **(D)** Immunofluorescence microscopy of HDAC5 in small intestine of conventional and germ-free mice. Arrowheads point to examples of HDAC5^+^ cells. Scale bars, 50 μm. Images are representative of 10 mice. **(E)** Immunofluorescence microscopy of HDAC5 in cells marked by staining with *Ulex europaeus* agglutinin 1 (UEA-I; arrowhead). Scale bars, 50 μm on the left and 10 μm on the right. **(F)** Barrier defense proteins are secreted by goblet cells and Paneth cells and assemble to form a mucus layer permeated with antimicrobial factors. **(G)** Immunoblot of IECs from the small intestines of wild-type and *Hdac5*^*-/-*^ mice with antibody detection of barrier defense proteins (LYZ1, ZG16, TFF3 and FCGBP). ACTIN was the loading control. **(H)** Band intensities of LYZ1, ZG16, TFF3 and FCGBP were measured by scanning densitometry and the relative densities were calculated using ACTIN as control. **(I)** MUC2 in wild-type and *Hdac5*^*-/-*^ IECs was measured by enzyme-linked immunosorbent assay (ELISA). MUC2 is not readily detected by immunoblot due to its high molecular weight and extensive glycosylation. **(J)** Immunofluorescence microscopy of barrier defense proteins in the small intestines of wild-type and *Hdac5*^*-/-*^ mice. Scale bars, 50 μm. qPCR, quantitative real-time PCR; IEC, intestinal epithelial cell; UEA-I, *Ulex europaeus* agglutinin I; LYZ1, lysozyme; ZG16, zymogen granule 16; TFF3, trefoil factor 3; FCGBP, F_c_ gamma binding protein; MUC2, mucin 2. Each experiment was performed at least twice; each bar graph data point represents one mouse; each immunoblot lane represents one mouse. Means ± SEM are plotted; *p < 0.05; **p < 0.01; by Student’s *t* test.

To investigate the function of HDAC5 in the intestine, we first identified the cell types that express it. In the small intestine, fluorescence microscopy revealed HDAC5 in a subset of IECs marked by *Ulex europaeus* agglutinin I (UEA-I), which labels secretory lineages including goblet cells in the villi and Paneth cells in the crypts (*14*) (Fig. 1, D and E). In the colon, HDAC5 localized to UEA-I^+^ epithelial cells consistent with goblet cells (fig. S1E).

Goblet and Paneth cells produce abundant proteins that are packaged into granules and secreted via exocytosis. Several of these proteins contribute to intestinal defense by forming a mucus barrier infused with antimicrobial factors. These include mucin 2 (MUC2), a glycoprotein that forms the structural scaffold of the mucus barrier (*1*); trefoil factor 3 (TFF3), which stabilizes mucin networks and initiates epithelial repair (*15, 16*); lysozyme (LYZ1), a bacteriolytic enzyme (*17*); zymogen granule protein 16 (ZG16), which aggregates bacteria (*4*); and Fc gamma binding protein (FCGBP), which inhibits bacterial attachment (*18*) (Fig. 1F). Levels of all five proteins were markedly reduced in the secretory epithelial cells of *Hdac5*^-/-^ mice (Fig. 1, G to J; fig. S1D; fig. S1, F to H).

In contrast, enterocyte expression of REG3G, an antimicrobial protein transcriptionally regulated by the microbiota, was mostly unaffected (fig. S1I). This suggests that HDAC5 selectively controls expression of barrier defense proteins whose expression is confined to secretory epithelial cells. Selective deletion of *Hdac5* in IECs (*Hdac5*^ΔIEC^ mice) abolished expression of the same set of proteins (fig. S1J). HDAC5 was also required for the expression of MUC2, ZG16, FCGBP and TFF3 in colonic goblet cells of *Hdac5*^*-/-*^ mice (note that LYZ1 is not expressed in the colon due to the absence of Paneth cells) (fig. S1K; fig. S1D). Thus, epithelial cell-intrinsic HDAC5 enables expression of a suite of barrier defense proteins in secretory epithelial cells of the small intestine and colon.

### HDAC5 maintains intestinal barrier integrity

The mucus layer and its embedded antimicrobial proteins maintain spatial segregation between the gut microbiota and the intestinal epithelium, preventing bacterial invasion of host tissue (*1, 19*). Given that HDAC5 promotes expression of multiple barrier defense proteins, we hypothesized that it is required to maintain this host-microbe separation. Fluorescence in situ hybridization revealed that bacteria were positioned closer to the epithelial surface in *Hdac5*^-/-^ than wild-type mice (Fig. 2, A and B). 16*S* rRNA gene sequencing further showed that loss of *Hdac5* altered microbial composition and reduced alpha diversity—hallmarks of a dysregulated intestinal environment (*20*)(Fig. 2, C and D; fig. S2, A and B).

**Figure 2.**
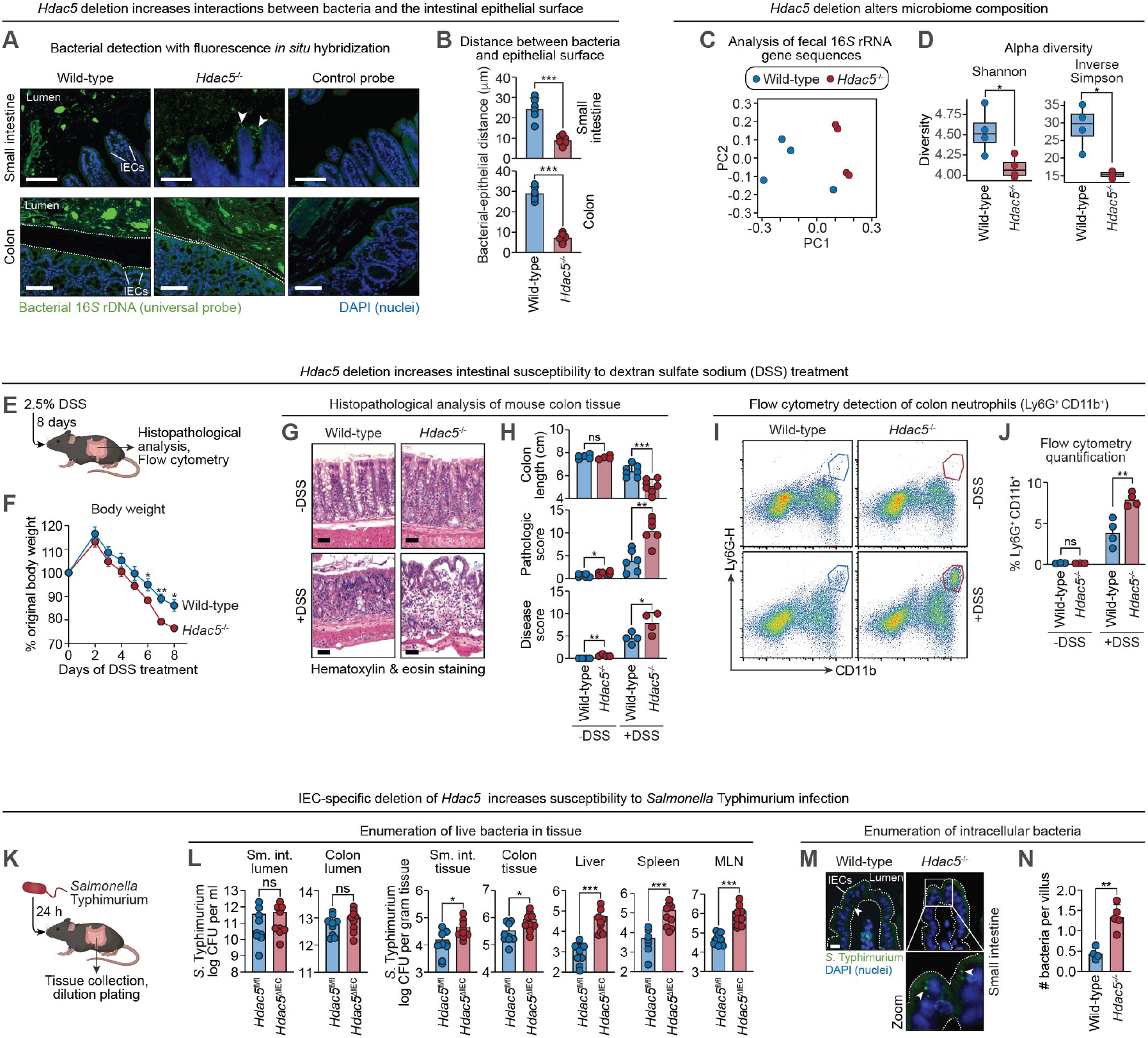
HDAC5 maintains intestinal barrier integrity. **(A)** Visualization of bacterial localization by fluorescence in situ hybridization and microscopy. Sections of mouse small intestine and colon were hybridized to a universal 16*S* rRNA gene probe and counterstained with 4’,6-diamidino-2-phenylindole (DAPI) to visualize nuclei. Sections are representative of 4 groups of littermates. Scale bars, 50 μm. **(B)** Measurement of distances between the villus tip to the nearest bacterial cells. Each data point is the average distance between a villus tip and the nearest bacteria in one mouse. Bacteria were identified on the basis of size, shape, and signal strength. 50-100 distances were measured per mouse, and 6 mice per genotype were analyzed. **(C)** Principal coordinate (PC) analysis of 16*S* rRNA sequencing of fecal samples from wild-type and *Hdac5*^-/-^ mice. The mice were littermates of heterozygous crosses that remained cohoused. **(D)** Alpha diversity measurements in fecal bacterial populations from wild-type and *Hdac5*^-/-^ mice. The mice were littermates of heterozygous crosses that remained cohoused. Each dot represents one mouse. **(E)** Experimental overview: mice were treated with 2.5% DSS for 8 days. Intestinal tissues were then analyzed for pathology and for changes to immune cells by flow cytometry. **(F)** Changes in body weight of wild-type and *Hdac5*^-/-^ mice after DSS treatment. N=3 mice/group. **(G)** Hematoxylin & eosin staining of colons from untreated and DSS-treated wild-type and *Hdac5*^-/-^ mice. Scale bars, 50 μm. **(H)** Colon length, histopathologic score and disease score of untreated and DSS-treated wild-type and *Hdac5*^-/-^ mice. **(I)** Representative flow cytometry plots of neutrophils (CD11b^+^Ly6G^+^) from the colon lamina propria of untreated and DSS-treated wild-type and *Hdac5*^-/-^ mice. **(J)** Frequencies of colon neutrophils from (I). N=3-4 mice/group. **(K)** Experimental overview: mice were orally inoculated with 5 × 10^9^ CFU of *Salmonella* Typhimurium. Tissues were collected after 24 hours and analyzed by dilution plating. **(L)** *S*. Typhimurium burdens in the lumen of small intestine (sm. int.) and colon, tissue of small intestine (sm. int.), colon, spleen, liver, and mesenteric lymph nodes (MLN) of *Hdac5*^fl/fl^ and *Hdac5*^ΔIEC^ littermates. N=9 mice/group. **(M)** Immunofluorescence microscopy of *S*. Typhimurium-GFP in small intestine epithelial cells. Sections were stained with a goat anti-GFP antibody and a donkey anti-goat IgG AlexaFluor^TM^ 488 secondary antibody. Examples of intracellular *S*. Typhimurium are indicated with arrowheads. Scale bars, 10 μm. **(N)** Enumeration of intracellular bacteria from (M). Fluorescent bacteria in each villus were counted and 50-100 villi were counted from each mouse. The average number of bacteria from each mouse was calculated and 4-5 mice were analyzed for each genotype. CFU, colony forming units. Each experiment was performed at least twice; each bar graph data point represents one mouse; each line graph data point represents the mean from three mice. Means ± SEM are plotted except for panel L, where geometric means ± SEM are plotted. *p < 0.05; **p < 0.01; ***p < 0.005; ns, not significant by Student’s *t* test.

*Hdac5*^-/-^ mice also exhibited heightened susceptibility to dextran sulfate sodium (DSS), which selectively damages the colonic epithelium (Fig. 2E). Following DSS treatment, *Hdac5*^-/-^ mice showed greater weight loss, more extensive epithelial damage, shortened colons, elevated histopathologic disease scores, and increased neutrophil infiltration (Fig. 2, F to J). In addition, *Hdac5* deficiency increased susceptibility to oral infection with *Salmonella enterica* serovar Typhimurium (*S*. Typhimurium). Although luminal bacterial loads were unchanged, *Hdac5*^ΔIEC^ mice had higher bacterial burdens in intestinal tissue, mesenteric lymph nodes, liver, and spleen than *Hdac5*^fl/fl^ controls (Fig. 2, K and L)—a phenotype also observed in global *Hdac5* knockout mice (fig. S2C). Moreover, there was more intracellular *S*. Typhimurium in IECs of *Hdac5*^-/-^ than wild-type mice, consistent with increased epithelial invasion (Fig. 2, M and N). Thus, HDAC5 is essential for optimal intestinal barrier function.

### HDAC5 stimulates mTORC1 signaling in intestinal epithelial cells

To identify the mechanism by which HDAC5 regulates barrier defense protein expression, we first tested whether HDAC5 acts through its canonical role as a transcriptional corepressor. Although this seemed unlikely given that HDAC5 enhances rather than represses protein production, we evaluated histone acetylation at target gene promoters. Histone deacetylases typically silence gene expression by removing acetyl groups from lysine residues on histone tails, thereby promoting chromatin condensation. Among these, acetylation of H3K9 and H3K27 is commonly associated with active transcription and is frequently used as a readout of HDAC activity (Fig. 3A). However, chromatin immunoprecipitation (ChIP)-qPCR showed no differences in acetylated H3K9 or H3K27 at barrier defense gene promoters in wild-type versus *Hdac5*^-/-^ epithelial cells (Fig. 3, B and C; fig. S3A). Furthermore, transcript levels remained unchanged despite reduced defense protein abundance in *Hdac5*^-/-^ mice (Fig. 3D; fig. S3B), arguing against a transcriptional mechanism. Supporting this, immunofluorescence microscopy revealed that HDAC5 was predominantly cytoplasmic and excluded from the nucleus in ileal and colonic epithelial cells (Fig. 3E; fig. S1E). This pattern was confirmed by biochemical fractionation of both mouse intestinal epithelial cells and HA-tagged HDAC5-expressing HT-29 cells (a human colonic epithelial cell line) (Fig. 3F; fig. S3C). Together, these findings suggested that HDAC5 regulates barrier defense protein expression via a non-canonical, cytoplasmic post-transcriptional mechanism.

**Figure 3.**
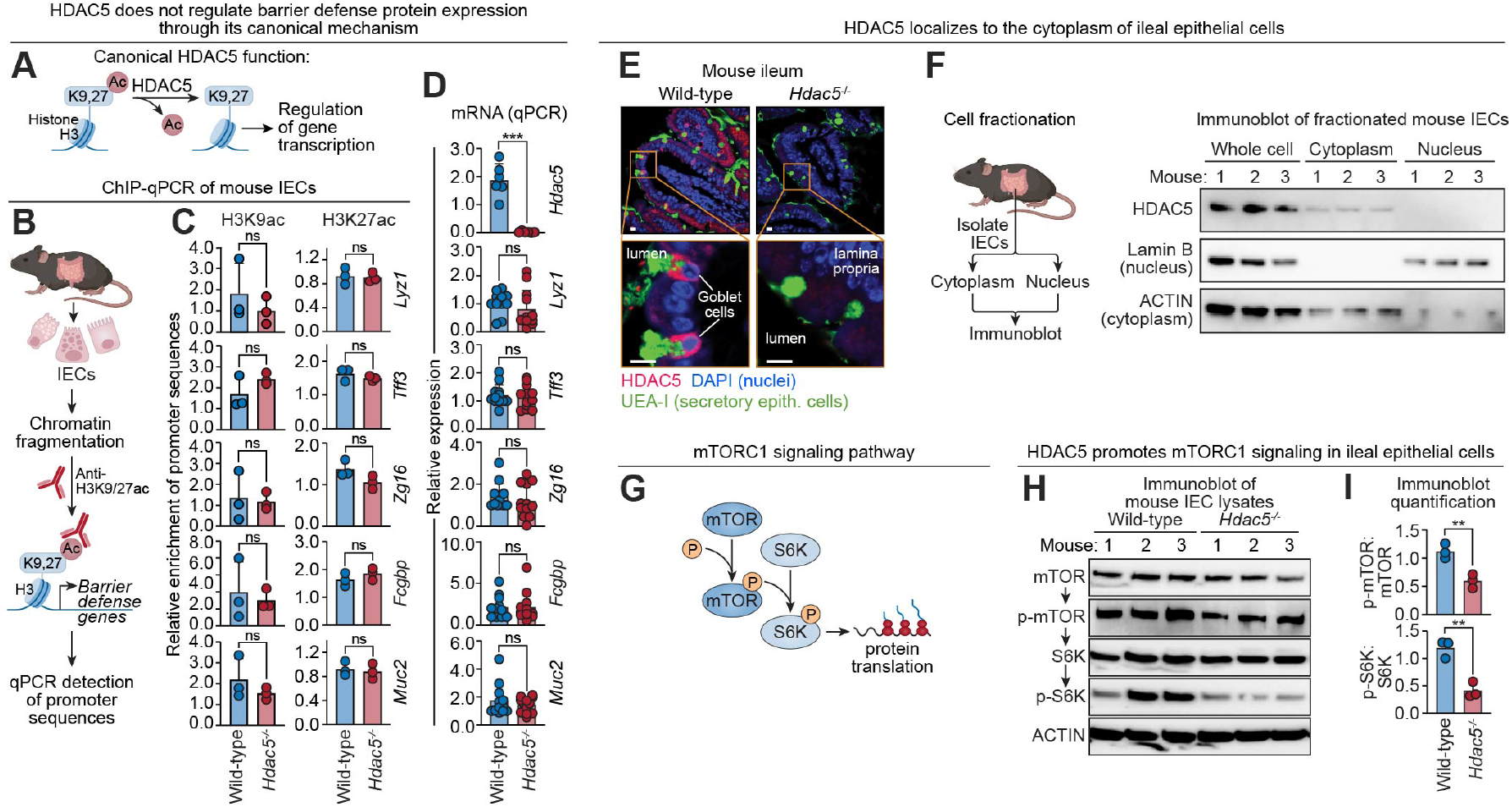
HDAC5 stimulates mTORC1 signaling in intestinal epithelial cells. **(A)** In its canonical function, HDAC5 deacetylates histone H3 at lysine 9 (K9) or lysine 27 (K27), leading to a closed chromatin configuration that suppresses target gene transcription. **(B)** Overview of ChIP-qPCR analysis of small intestine IECs. Histone acetylation at the promoters of barrier defense genes was compared in wild-type and *Hdac5*^-/-^ mice. **(C)** Measurement of H3K9ac and H3K27 marks at the promoters of barrier defense genes in IECs recovered from wild-type and *Hdac5*^*-/-*^ mice, using the scheme in (B). N=3 mice per group. **(D)** qPCR of barrier defense gene transcripts in IECs from wild-type and *Hdac5*^*-/-*^ mice. N=12 mice/group (N=6 for analysis of *Hdac5*). **(E)** Immunofluorescence microscopy of HDAC5 in the small intestines of wild-type and *Hdac5*^*-/-*^ mice indicates that HDAC5 is mostly excluded from the nuclei of secretory epithelial cells. Scale bars, 10 μm. **(F)** Small intestine epithelial cells recovered from wild-type mice were separated by centrifugation into cytoplasm- and nucleus-containing fractions. The fractions were immunoblotted and HDAC5 was detected alongside Lamin B (a marker for nuclei) and ACTIN (a marker for cytoplasm). **(G)** Regulation of protein translation by the mTORC1 signaling pathway. **(H)** Immunoblot detection of mTORC1 signaling pathway components in IEC lysates from wild-type and *Hdac5*^*-/-*^ mice. **(I)** Band intensities from (H) were measured by scanning densitometry and protein ratios were calculated. IECs, intestinal epithelial cells; UEA-I, *Ulex europaeus* agglutinin 1. Each experiment was performed at least twice; each bar graph data point represents one mouse; each immunoblot lane represents one mouse. Means ± SEM are plotted; **p < 0.01; ***p < 0.001; ns, not significant by Student’s *t* test.

We next sought to define the mechanism by which cytoplasmic HDAC5 enhances protein production in secretory epithelial cells. One well-established means of increasing protein output is by promoting translation of mRNAs. In many cells, this process is regulated by the mechanistic target of rapamycin (mTOR), a serine/threonine kinase that integrates signals of energy status and other environmental cues (*21*). mTOR regulates energetically costly processes such as translation by activating S6 kinase (S6K), which phosphorylates components of the translational machinery (*22, 23*) (Fig. 3G). mTOR operates in two distinct multiprotein complexes, mTORC1 and mTORC2, with mTORC1 being the primary driver of protein synthesis (*21*). Notably, mTORC1 signaling has been reported in Paneth cells, where it links intestinal stem cell function to energy availability (*24*). Importantly, prior studies in the optic nerve and stomach have shown that cytoplasmic HDAC5 can activate mTOR signaling in those tissues (*25, 26*), raising the possibility that HDAC5 might enhance translation in IECs by promoting mTORC1 activity.

To test whether HDAC5 modulates mTORC1 signaling in IECs, we performed immunoblot analysis of key pathway components in primary IECs harvested from mouse intestines. Levels of phosphorylated mTOR (p-mTOR) and phosphorylated S6 kinase (p-S6K) were reduced in ileal and colonic epithelial cells from *Hdac5*^-/-^ mice compared to wild-type controls (Fig. 3, H and I; fig. S3, D and E). These findings indicate that HDAC5 promotes mTORC1 signaling and suggest that it may enhance post-transcriptional expression of barrier defense proteins through the mTORC1 pathway.

### HDAC5 promotes barrier defense protein translation through mTORC1

We tested whether HDAC5 promotes barrier defense protein expression through mTORC1 by treating mice with rapamycin, an inhibitor of mTORC1 signaling and downstream protein translation (*27, 28*) (Fig. 4, A and B). Rapamycin treatment reduced levels of p-mTOR and p-S6K in IECs from both the ileum and colon (Fig. 4, C and D; fig. S4, A and B), while also reducing LYZ, TFF3, ZG16, and FCGBP levels (Fig. 4, C and D; fig. S4, A and B). We observed similar results in LS513 cells, a human IEC line, where rapamycin treatment decreased levels of LYZ1, ZG16, and FCGBP without reducing their transcript abundance (fig. S4, C to F). These data indicate that mTORC1 signaling enhances post-transcriptional production of barrier defense proteins in IECs.

**Figure 4.**
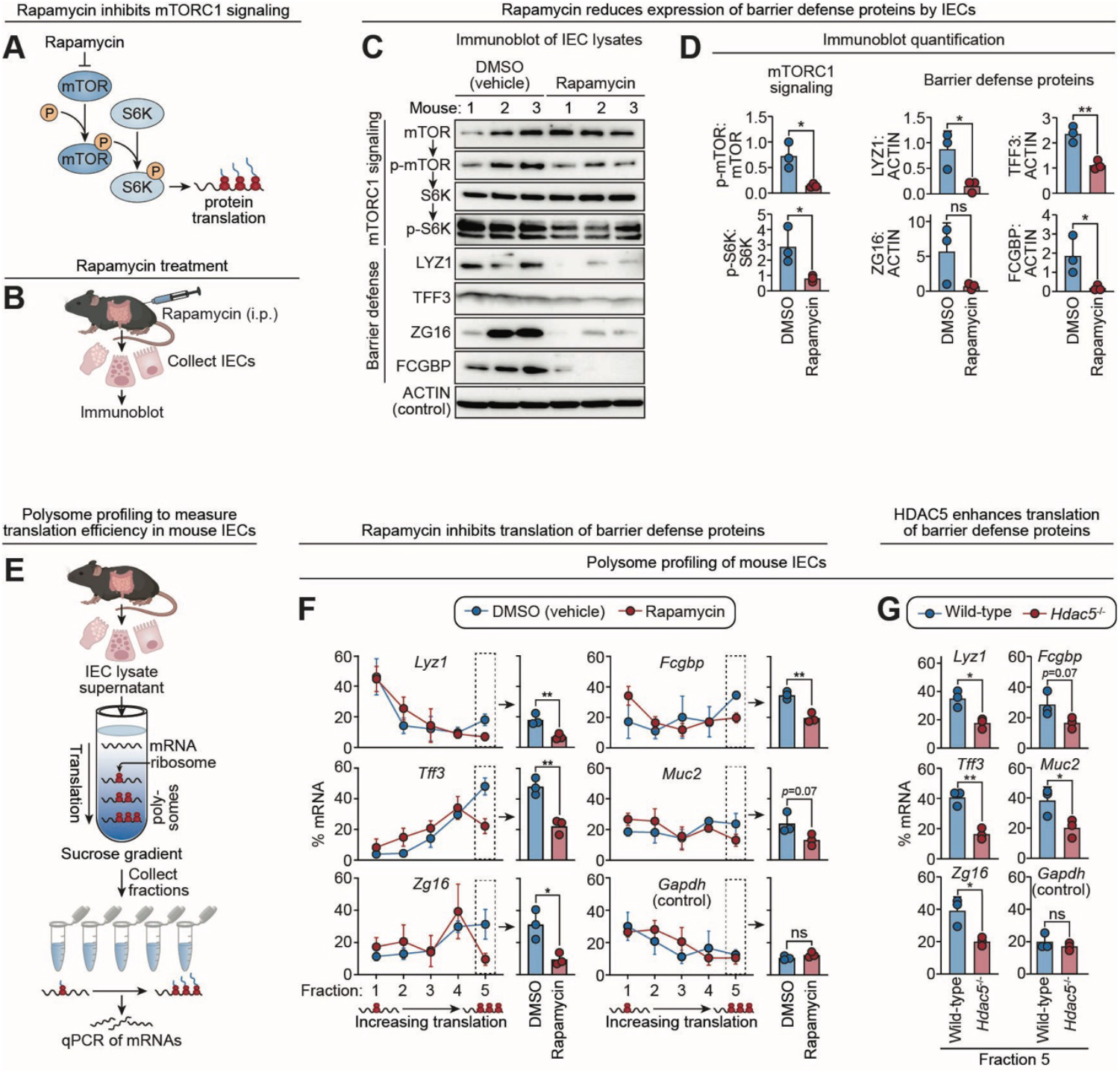
HDAC5 promotes barrier defense protein translation through mTORC1. **(A)** Rapamycin reduces protein translation by inhibiting mTORC1 activity (*21*). **(B)** Experimental scheme for (C) and (D). Mice were injected with rapamycin (i.p.=intraperitoneal route) on each of three days and small intestine IECs were analyzed by immunoblot. **(C)** Immunoblot of IEC lysates from mice treated with rapamycin or DMSO (vehicle control) with detection of mTORC1 signaling pathway components and barrier defense proteins. ACTIN was the loading control. Note that MUC2 is not readily detected by immunoblot due to its large size. **(D)** Band intensities from (C) were measured by scanning densitometry and protein ratios were calculated. **(E)** Overview of polysome profiling assays to analyze barrier defense protein translation in small intestine epithelial cells. IEC lysates were loaded onto a linear sucrose gradient and separated by ultracentrifugation. mRNAs localize along this gradient according to the number of bound ribosomes, which correlate with translation rate. RNA was isolated from individual gradient fractions and mRNAs encoding barrier defense proteins were measured by qPCR. **(F)** Polysome profiling of barrier defense gene transcripts in rapamycin-treated and DMSO (vehicle)-treated mice. The percentage of transcripts from each gradient fraction are shown by line graphs on the left. Fraction 5 contains mRNA associated with the highest number of ribosomes, indicating a high rate of translation. *Gapdh* transcripts were analyzed as a control. **(G)** Polysome profiling of barrier defense gene transcripts in wild-type and *Hdac5*^-/-^ mice. mRNA percentages from fraction 5 are shown, representing the highest translation rates. The complete gradient line plots for each mRNA are shown in Figure S4G. IEC, intestinal epithelial cells; UEA-I, *Ulex europaeus* agglutinin I; LYZ1, lysozyme; ZG16, zymogen granule 16; TFF3, trefoil factor 3; FCGBP, Fc gamma binding protein. Each experiment was performed at least twice; each bar graph data point represents one mouse; each line graph data point represents the mean from three mice; each immunoblot lane represents one mouse. Means ± SEM are plotted; *p < 0.05; **p < 0.01; ns, not significant by Student’s *t* test.

Regulation of translational efficiency is a key mechanism for controlling protein abundance. Polysome profiling assesses translational efficiency by separating mRNAs based on the number of bound ribosomes (*29*). Poorly translated transcripts associate with few ribosomes, while actively translated ones are found in high-density polysome fractions. To test whether mTORC1 regulates translation of intestinal barrier defense proteins, we performed polysome profiling on primary IECs harvested from rapamycin-treated mice (Fig. 4E). Sucrose gradient fractionation followed by qPCR revealed reduced levels of barrier defense transcripts in high-density fractions, indicating decreased translation efficiency (Fig. 4F). Similarly, polysome profiling of *Hdac5*^-/-^ IECs, in which mTORC1 signaling is impaired (Fig. 4, C and D), showed reduced polysome association of these transcripts (Fig. 4G; fig. S4G). Notably, we did not observe this effect for *Gapdh* transcripts, indicating that HDAC5-mTORC1 signaling selectively regulates translation of barrier defense mRNAs rather than globally suppressing protein synthesis. Together, these results suggest that HDAC5 promotes barrier defense protein production by enhancing mTORC1-dependent translation.

### HDAC5 activates mTORC1 by deacetylating 14-3-3 to sequester Raptor

We next sought to define the mechanism by which HDAC5 activates mTORC1 signaling. To identify the HDAC5 substrate in the intestinal epithelial cell cytoplasm, we conducted an unbiased screen for HDAC5-interacting proteins. We expressed HA-tagged HDAC5 in *HDAC5*^-/-^ human HT-29 colonic epithelial cells and performed anti-HA immunoprecipitation to capture interacting protein partners (Fig. 5, A and B). Liquid chromatography–tandem mass spectrometry (LC–MS/MS) identified multiple isoforms of 14-3-3, with 14-3-3ε being the most abundant (Fig. 5C; Table S1). This interaction was confirmed in mouse IECs by reciprocal co-immunoprecipitation with anti–14-3-3 and anti–HDAC5 antibodies and is consistent with prior reports of HDAC association with 14-3-3 proteins in non-epithelial cells (*30*). These data thus support a physical association between HDAC5 and 14-3-3 proteins (Fig. 5D).

**Figure 5.**
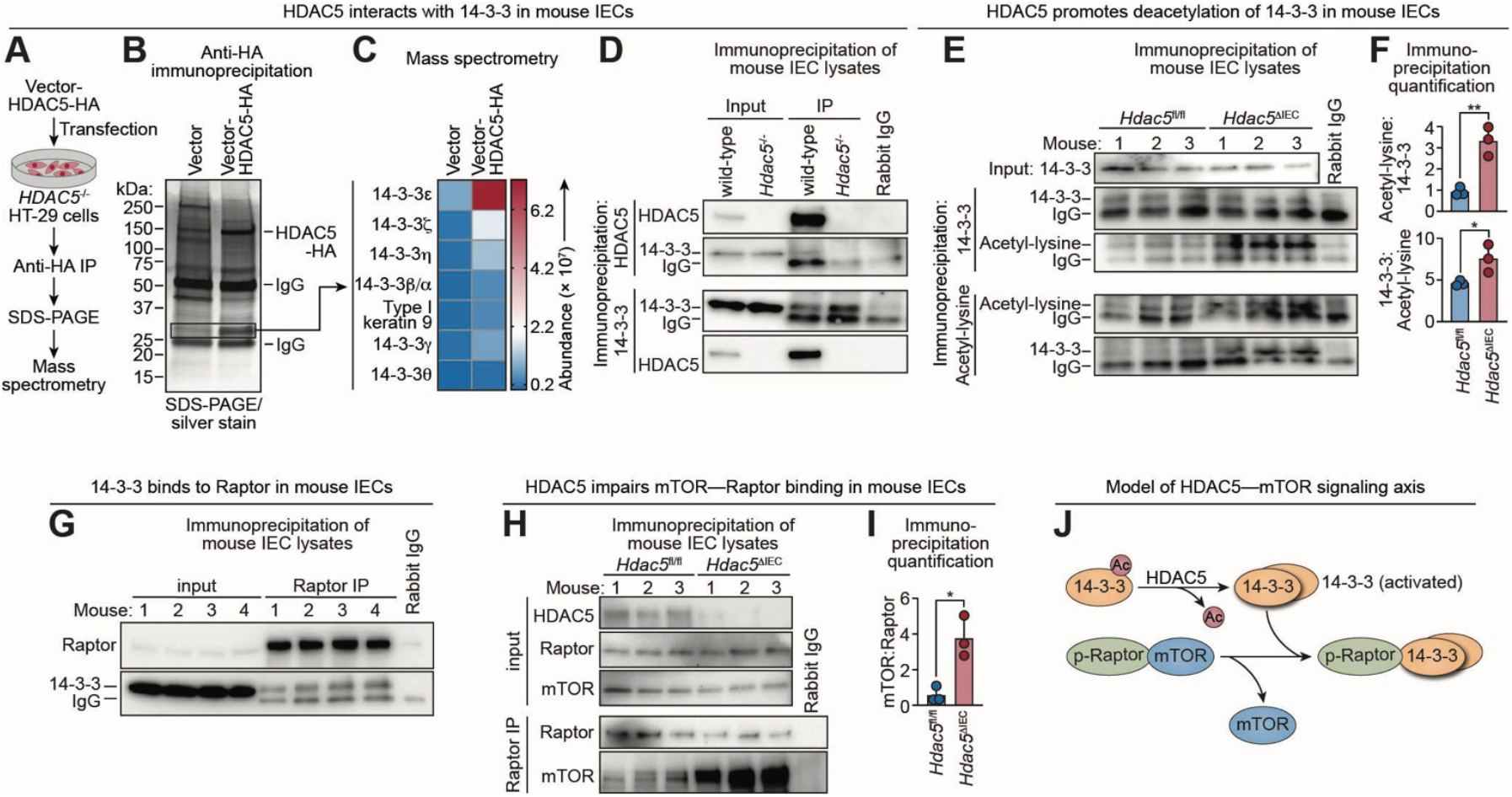
HDAC5 activates mTORC1 by deacetylating 14-3-3 to sequester Raptor. **(A)** Identification of HDAC5 interacting partners/substrates. Human HDAC5 was fused to a hemagglutinin (HA) tag and the vector-HDAC5-HA construct was transfected into HT-29 human colorectal carcinoma cells. Cell lysates were immunoprecipitated with an anti-HA antibody and separated by SDS-PAGE. HDAC5-interacting proteins were identified by liquid chromatography–tandem mass spectrometry (LC–MS/MS). **(B)** Silver stained SDS-PAGE gel of anti-HA immunoprecipitation pellets from the experiment outlined in (A). The indicated bands were analyzed by LC–MS/MS. **(C)** LC–MS/MS identification of proteins in the SDS-PAGE bands shown in (B). **(D)** Detection of HDAC5 interaction with 14-3-3. HDAC5 or 14-3-3 was immunoprecipitated from small intestine IEC lysates and blotted, with detection of HDAC5 and 14-3-3. Each lane represents an immunoprecipitation on pooled lysates from three mice. **(E)** Detection of acetylated 14-3-3. 14-3-3 or acetyl-lysine were immunoprecipitated from small intestine IEC lysates and blotted, with detection of both 14-3-3 and acetyl-lysine. **(F)** Band intensities in (E) were measured by scanning densitometry and protein ratios were calculated. **(G)** Small intestine IECs were recovered from wild-type mice, lysed, immunoprecipitated with anti-Raptor antibody, and blotted alongside total lysate (input). The precipitated proteins were blotted and probed with anti-Raptor and anti-14-3-3. **(H)** Small intestine IECs were recovered from *Hdac5*^fl/fl^ and *Hdac5*^ΔIEC^ mice, lysed, immunoprecipitated with anti-Raptor antibody, and blotted alongside total lysate (input). The blots were probed with anti-Raptor or anti-mTOR antibodies. **(I)** Band intensities from (H) were measured by scanning densitometry and protein ratios were calculated. **(J)** Model showing how HDAC5 promotes mTORC1 activity by regulating Raptor binding to mTOR. HDAC5 deacetylates 14-3-3, which becomes active via self-association, binds phosphorylated Raptor, and releases mTOR to engage in downstream signaling pathways. HA, hemagglutinin tag; IP, immunoprecipitation; IEC, intestinal epithelial cells; Each experiment was performed at least twice; each bar graph data point represents one mouse; each immunoblot lane represents one mouse except in (D), where lysates from multiple mice were pooled. Means ± SEM are plotted; *p < 0.05; **p < 0.01; ns, not significant by Student’s *t* test.

14-3-3 proteins are a family of conserved signaling adaptors that bind phosphorylated proteins, and this binding depends on 14-3-3 dimerization (*31*). Acetylation disrupts 14-3-3 self-association, whereas deacetylation promotes it (*32*). To test whether HDAC5 deacetylates 14-3-3 we assessed 14-3-3 acetylation in IECs recovered from wild-type and *Hdac5*^-/-^ mice. Immunoprecipitation with anti–14-3-3 or anti–acetyl-lysine antibodies followed by immunoblotting revealed increased 14-3-3 acetylation in *Hdac5*^-/-^ IECs (Fig. 5, E and F). LC-MS/MS analysis of proteins associated with HA-tagged 14-3-3ε further revealed greater 14-3-3 self-association in wild-type than *HDAC5*^-/-^ HT-29 cells (fig. S5, A to C; Table S2), consistent with the known inhibitory effect of acetylation. Together, these findings indicate that HDAC5 promotes the deacetylation of 14-3-3 in IECs and suggest that deacetylation enables 14-3-3 self-association, potentially enhancing its ability to engage phosphorylated targets.

Phosphorylated Raptor (p-Raptor) is a known phosphoprotein binding target of 14-3-3 that is part of the mTORC1 signaling pathway (*33*). Although Raptor promotes mTORC1 activity in some cellular contexts, AMPK-mediated phosphorylation converts it to an mTORC1 suppressor under conditions of energy stress (*33*). With this mechanism in mind, we hypothesized that HDAC5 enhances mTORC1 signaling by deacetylating 14-3-3, thus promoting its self-association and facilitating the sequestration of inhibitory p-Raptor. To test this, we immunoprecipitated Raptor from wild-type small intestinal epithelial cell lysates and detected associated 14-3-3, confirming the interaction of these proteins (Fig. 5G). Furthermore, more mTOR co-immunoprecipitated with Raptor in IECs from *Hdac5*^ΔIEC^ mice than from *Hdac5*^fl/fl^ controls (Fig. 5, H and I), consistent with impaired Raptor– 14-3-3 interaction and increased Raptor–mTOR association in the absence of HDAC5. These findings support a signaling pathway in which epithelial HDAC5 deacetylates 14-3-3, enabling it to sequester Raptor. This relieves mTORC1 inhibition and enhances protein translation (Fig. 5J).

### The microbiota enhances translation of barrier defense proteins through the HDAC5—mTORC1 pathway

Since the microbiota induces HDAC5 expression (Fig. 1 B and C) and HDAC5 promotes mTORC1-dependent translation of barrier defense proteins (Fig. 4), we next asked whether the microbiota enhances protein translation through the HDAC5—mTORC1 pathway. Specifically, we tested whether microbial colonization promotes 14-3-3 deacetylation, activates mTORC1, and increases translation of barrier defense proteins in an HDAC5-dependent manner. To do so, we compared primary IECs from wild-type germ-free and conventional mice, as well as from conventional *Hdac5*^ΔIEC^ mice. 14-3-3 acetylation was elevated in both germ-free wild-type and conventional *Hdac5*^ΔIEC^ mice relative to conventional controls (Fig. 6, A and B), consistent with HDAC5-mediated deacetylation downstream of the microbiota. Immunoblotting revealed reduced levels of p-S6K in both groups (Fig. 6C; fig. S6, A and B), suggesting that the microbiota activates mTORC1 signaling through HDAC5. Finally, polysome profiling showed decreased association of barrier defense transcripts with high-density fractions in both groups, indicating reduced translational efficiency in the absence of either microbial colonization or epithelial HDAC5 (Fig. 6D). Together, these findings support a model in which the microbiota enhances translation of barrier defense proteins by inducing HDAC5, which deacetylates 14-3-3 to activate mTORC1 signaling (Fig. 6E).

**Figure 6.**
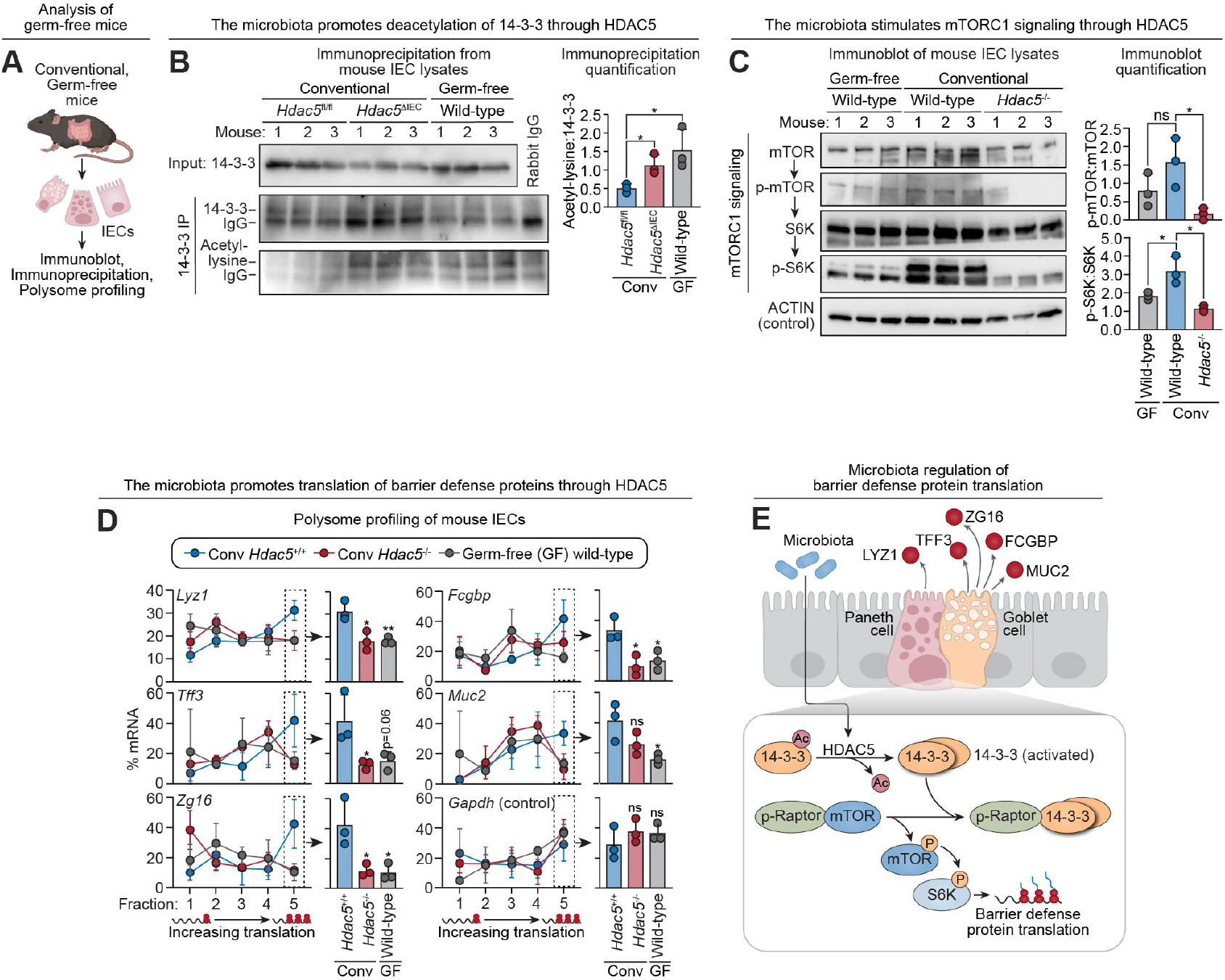
The microbiota enhances translation of barrier defense proteins through the HDAC5—mTORC1 pathway. **(A)** Experimental scheme for (B) and (C). IECs were recovered from conventional or germ-free mice and analyzed by immunoblot or immunoprecipitation. **(B)** Immunoprecipitations to detect acetylation of 14-3-3. 14-3-3 was immunoprecipitated from small intestine IEC lysates and blotted, with detection of 14-3-3 and acetyl-lysine. Total lysate input was analyzed as a control. Band intensities were measured by scanning densitometry and protein ratios were calculated. **(C)** Immunoblot of small intestine IEC lysates from conventional mice (wild-type, *Hdac5*^*-/-*^) and germ free (wild-type) mice, with detection of mTORC1 signaling pathway components. ACTIN was the loading control. Band intensities were measured by scanning densitometry and protein ratios were calculated. **(D)** Polysome profiling of small intestine IECs from conventional mice (wild-type, *Hdac5*^-/-^) and germ-free (wild-type) mice. The percentage of barrier defense protein mRNAs from each gradient fraction are shown by line graphs on the left. Fraction 5 contains mRNA associated with the highest number of ribosomes, indicating a high rate of translation. *Gapdh* was used as a control. **(E)** Proposed model of microbiota-driven regulation of barrier defense protein translation. The gut microbiota induces expression of HDAC5, which promotes deacetylation of the regulatory protein 14-3-3. Deacetylated 14-3-3 binds phosphorylated Raptor, disrupting the mTOR-Raptor complex and enhancing mTOR activity. Activated mTOR signaling promotes translation of barrier defense proteins, which in turn regulate microbial interactions with the intestinal surface. IEC, intestinal epithelial cells; IP, immunoprecipitation; Conv, conventional; GF, germ-free. Each experiment was performed at least twice; each bar graph data point represents one mouse; each line graph data point represents the mean from three mice; each immunoblot lane represents one mouse. Means ± SEM are plotted; *p < 0.05; **p < 0.01; ns, not significant by Student’s *t* test.

Our finding that the microbiota enhances translation of mucus barrier proteins was unexpected, as prior studies of proteins such as LYZ1 have not reported substantial differences in their tissue abundance between germ-free or antibiotic-treated mice and conventional controls (*10*). Our own analyses confirm this observation (fig. S7, A to C). This apparent paradox is explained by earlier reports that bacteria induce protein secretion by secretory IECs (*34*). Consistent with this, we observed reduced release of barrier defense proteins into the intestinal lumen in antibiotic treated mice (fig. S8A), and increased release upon microbial stimulation of small intestinal organoids (fig. S8, B to D). Thus, steady-state epithelial protein levels reflect the balance between synthesis and export, both of which are microbiota-dependent. In the absence of the microbiota, reduced secretion permits intracellular accumulation of barrier defense proteins despite diminished translation, obscuring changes in production rate (fig. S8E). Coordinated regulation of translation and secretion may enable the epithelium to deploy barrier defenses efficiently in response to microbial cues and could explain why microbiota-dependent control of protein synthesis has previously gone unrecognized.

## Discussion

We have identified a pathway by which the gut microbiota enhances translation of key barrier defense proteins, including lysozyme and mucin 2. Microbial colonization induces secretory epithelial cell expression of HDAC5, which deacetylates 14-3-3 proteins. This enables 14-3-3 to sequester Raptor, relieving inhibition of mTORC1 and stimulating translation of barrier defense proteins. These proteins are secreted apically in response to microbial cues, assembling into a mucus barrier permeated with antimicrobial effectors that restrict bacterial access to host tissue. When considered alongside prior findings that the microbiota induces transcription of a distinct set of antimicrobial genes (*6-8*), our findings suggest that the microbiota orchestrates barrier immunity through both transcriptional and translational control.

Previous studies have shown that bacterial metabolites influence mTOR activity in immune cells and colonocytes (*35, 36*). Further, bacterial pathogen effector proteins can modulate mTORC1 activity to liberate host nutrients for pathogen consumption (*37*). Here we have uncovered a distinct mechanism of bacterial modulation of mTORC1 signaling in secretory IECs. Specifically, we identify HDAC5 as a critical intermediary that links microbial colonization to mTORC1 activation. As a key metabolic gatekeeper, mTORC1 integrates signals of nutrient availability to regulate energy-intensive processes such as protein synthesis. This regulation is likely to be especially important in secretory IECs, where production of large quantities of mucus infused with abundant antimicrobial proteins requires substantial translational capacity. Integrating microbial and nutrient signals through mTOR could ensure that barrier immunity is activated only when both energetic conditions and microbial threats justify the metabolic investment.

Several key questions remain. First, it is not yet clear whether HDAC5 deacetylates 14-3-3 directly or whether it acts indirectly through interaction with other deacetylases, such as HDAC3. This possibility is suggested by prior structural and biochemical studies showing that class IIa HDACs (including HDAC5) are inefficient deacetylases on acetyl-lysine substrates due to a catalytic tyrosine-to-histidine substitution that impairs activity (*38*). Second, it is unclear how the HDAC5–mTORC1 axis intersects with other microbe-regulated pathways controlling mucus secretion, such as inflammasome signaling and autophagy (*39, 40*). Third, during pathogen infection, lysozyme is rerouted into the secretory autophagy pathway (*41*), and it is unknown how this alternate trafficking integrates with microbiota-enhanced translation. Finally, the upstream signals by which the microbiota induces *Hdac5* expression remain to be defined.

Our data suggest that disrupting the HDAC5—mTORC1 signaling axis could weaken immunity and increase susceptibility to inflammation and infection. Although *HDAC5* has not been linked to inflammatory bowel disease (IBD) in genetic studies, histone deacetylases more broadly have been implicated in disease pathogenesis through epigenetic and functional analyses. In particular, pan-HDAC inhibitors are protective in mouse models of IBD, suggesting the therapeutic potential of targeting HDAC activity. These findings position HDAC5 as a potential target for microbiome- or drug-based interventions aimed at restoring barrier integrity.

## Acknowledgements

We thank E. Olson (UT Southwestern) for the *Hdac5*^-/-^ mice, C. Salinas and M. Johnson for assistance with germ-free mouse experiments, J. Otto (UT Southwestern Proteomics Core) for assistance with mass spectrometry, and B. Zhang (UT Southwestern Microbiome Research Laboratory) for assistance with 16*S* rRNA gene sequencing.

## Author contributions

Y.L., Z.K., and L.V.H. designed research. Y.L., R.W., X.M., J.Z., J.M., Y.D., B.H., K.A.R., C.L.B., C.D., L.W., A.L. and P.R. performed research. Y.L., A.L., P.R. and Z.K. analyzed data. Y.L., Z.K., and L.V.H. wrote the paper.

## Data and materials availability

16*S* rRNA gene sequencing data are available from the Gene Expression Omnibus (GEO) repository under accession number GSE304136. All other data are available in the main text or the supplementary materials.

## Supplementary Materials

**Materials and Methods**

**Figures S1 to S8**

**Tables S1-S5**

