## Supplementary materials for "Translational control of innate barrier defense by the gut microbiota"

**This PDF file includes:**

Materials and Methods
Figures S1 to S8
Tables S1-S5

### Materials and Methods

#### Mice

Wild-type C57BL/6 and *Hdac5*$^{-/-}$ mice (*42*) were bred and maintained under specific pathogen–free (SPF) conditions at the University of Texas Southwestern Medical Center. *Hdac5*$^{fl/fl}$ mice (*43*) were crossed with Villin-Cre transgenic mice (*44*) to generate intestinal epithelial cell–specific knockouts (*Hdac5*$^{\Delta IEC}$). All mice were maintained on standard chow (LabDiet 5KA1) containing 22% protein, 16% fat, and 62% carbohydrates, provided ad libitum. Germ-free C57BL/6 mice were housed and bred in the gnotobiotic facility at UT Southwestern, as described previously (*6*). All experiments were conducted using 8- to 12-week-old male and female mice. Animals were housed on a 12-hour light/12-hour dark cycle. Mice were euthanized by overdose of inhaled isoflurane followed by cervical dislocation. All procedures were approved by the Institutional Animal Care and Use Committee (IACUC) at the University of Texas Southwestern Medical Center.

#### Antibiotic treatment of mice

Conventionally-raised C57BL/6 mice were maintained on drinking water containing 1 mg/ml neomycin, 1 mg/ml gentamycin, 1 mg/ml metronidazole, 1 mg/ml streptomycin, and 0.5 mg/ml vancomycin for at least two weeks. Microbiota depletion was verified by aerobic and anaerobic culture of fecal pellets.

#### Isolation of mouse intestinal epithelial cells

Mouse ileum (distal small intestine) and colon tissues were washed with ice-cold PBS, opened longitudinally, and cut into ~1 cm segments. Tissue pieces were placed in 20 ml of cold PBS and washed three times with vigorous shaking. Once the supernatant appeared clear, the tissue was transferred to a fresh tube containing 20 ml of 10 mM ethylenediaminetetraacetic acid (EDTA) in PBS and rotated at 4°C for 15 minutes. Intestinal epithelial cells were released by vigorous shaking and collected by centrifugation at 300 × *g* for 3 minutes. The cell pellets were washed twice with PBS and used directly in downstream assays.

**Quantitative real-time PCR (qPCR)**

Complementary DNA (cDNA) was generated from purified RNA using M-MLV Reverse Transcriptase (Fisher, 28025021). Quantitative real-time PCR was conducted on a QuantStudio 7 Flex Real-Time PCR System (Applied Biosystems) with Platinum SYBR Green RT-qPCR SuperMix (Fisher, 11733046). Transcript levels were quantified using the comparative Ct (ΔΔCt) method (*45*) and normalized to *Gapdh* expression. Primer sequences used for qPCR are provided in Table S3.

**Immunoblot**

Epithelial cells isolated from mouse intestines were resuspended in 250 µl of T-PER buffer (Thermo Scientific) supplemented with protease and phosphatase inhibitor cocktail (Thermo Scientific, 78440) and deacetylase inhibitor cocktail (ApexBio, K1017). Suspensions were incubated on ice for 10 minutes and centrifuged at 20,000 × *g* for 10 minutes at 4°C to generate clarified cell lysates. Proteins from human colorectal adenocarcinoma cell lines (LS513 and HT-29) were extracted using the same procedure.

Protein concentrations were measured using a NanoDrop One spectrophotometer (Thermo Scientific) and normalized across samples. Equal amounts of protein were separated by SDS-PAGE using 4–20% gradient gels and transferred to PVDF membranes. Membranes were blocked with 5% blotting-grade blocker (Bio-Rad) in TBS-T buffer (Tris-buffered saline with 0.1% Tween-20) for 1 hour at room temperature, then incubated sequentially with primary and HRP-conjugated secondary antibodies (listed in Table S5). Protein bands were visualized using a Bio-Rad ChemiDoc imaging system.

**Immunofluorescence microscopy**

Mouse ileum and colon tissues were flushed with PBS and fixed overnight at 4°C in Bouin's fixative (Fisher Scientific, 112016), then embedded in paraffin. For immunostaining of proteins secreted into the intestinal lumen, unflushed small intestine and colon tissues were fixed in methacarn solution (60% methanol, 30% chloroform, 10% glacial acetic acid) and similarly embedded in paraffin.

Tissue sections were deparaffinized by two 10-minute washes in xylene, followed by two 3-minute washes in 100% ethanol, two washes in 95% ethanol, two washes in 70% ethanol, and a final rinse in deionized water for

5 minutes. Antigen retrieval was performed by boiling sections in 10 mM sodium citrate buffer (pH 6.0) for 20 minutes, followed by washing in 1× PBS. Sections were blocked sequentially in 2% BSA in PBS for 15 minutes, then in 1% BSA with 0.3% Triton X-100 in PBS for 15 minutes. After additional PBS washes, sections were incubated with primary antibodies (listed in Table S5) overnight at 4°C. After three washes in PBS-T buffer (0.1% Tween-20 in PBS), sections were incubated with fluorescent dye–conjugated secondary antibodies (Table S5) or UEA-I (*Ulex europaeus* agglutinin I) Lectin (FITC) (GeneTex, GTX01512) for 1 hour at room temperature. Nuclei were counterstained with DAPI, and images were acquired using a Zeiss AxioImager M1 microscope.

**Mucin 2 (MUC2) enzyme-linked immunosorbent assay (ELISA)**

Ileal tissues were collected from wild-type and *Hdac5*$^{-/-}$ mice and rinsed and homogenized in ice-cold PBS. Lysates were frozen and thawed for two cycles, and the homogenates were centrifuged for 5 minutes at 5000 × *g*. The supernatants were recovered and MUC2 was measured using a mouse mucin 2 ELISA kit (MyBioSource, MBS2886327).

**Fluorescence in situ hybridization (FISH)**

Fluorescence in situ hybridization (FISH) was used to localize bacteria in the small intestine and colon using a universal 16*S* rRNA gene probe (Table S1). Ileal and colonic tissue fragments were fixed in methacarn (60% methanol, 30% chloroform, 10% acetic acid) and washed three times in 70% ethanol. The tissues were embedded in paraffin and sections were cut by the UT Southwestern Histopathology Core. Sections were deparaffinized in xylene (2 × 10 min) and rehydrated through graded ethanol (100%, 95%, 90%, 70%) to water (2 × 5 min per step). Hybridization buffer (0.9 M NaCl, 20 mM Tris-HCl pH 7.2, 0.1% SDS) was pre-warmed to 56°C, and the probe (sequence listed in Table S3) was applied at 10 nM with a glass coverslip. Hybridization was performed overnight at 50°C in a humidified chamber alongside a non-specific control probe (sequence listed in Table S3). Slides were washed (3 × 10 min) in buffer (0.9 M NaCl, 20 mM Tris-HCl, pH 7.2), mounted with DAPI Fluoromount-G (SouthernBiotech), and imaged using a Zeiss AxioImager M1 microscope. Images were analyzed with ImageJ. The distance between villus tips and the nearest bacteria was measured in pixels and converted to microns (μm)(for 20X images, 1 pixel=0.75 μm; for 40X images, 1 pixel=0.375 μm).

**16*S* rRNA gene sequencing and data analysis**

DNA was isolated from freshly collected fecal pellets using the FastDNA Spin Kit (MP Biomedicals 116560-200) in combination with a FastPrep-24 5G Homogenizer. To generate sequencing libraries, the V3–V4 variable regions of the 16*S* rRNA gene were amplified using HotStarTaq Plus Master Mix (Qiagen) and region-specific primers (Table S3). Amplicons were sequenced on an Illumina MiSeq platform according to the manufacturer's instructions. Reads were clustered into operational taxonomic units (OTUs) at 97% sequence identity (3% divergence). Taxonomic classification of OTUs was performed using BLASTn against a custom-curated database combining RDPII and NCBI sequences. Raw OTU abundance data were first imported and formatted using the R package phyloseq. Relative abundance tables were converted into phyloseq-standard OTU tables. OTUs with zero total abundance across all samples were removed. Alpha diversity metrics (Shannon, Inverse Simpson, Observed species, Chao1, ACE, Simpson, and Fisher indices) were statistically tested using both Wilcoxon rank-sum tests and independent t-tests to assess non-parametric and parametric differences.

**Treatment of mice with dextran sulfate sodium (DSS)**

Wild-type and *Hdac5*$^{-/-}$ mice were administered 2.5% DSS (m.w. 36,000–50,000; MP Chemicals) in drinking water ad libitum for up to 8 days, with daily monitoring of body weight and fluid intake. DSS consumption was comparable across groups. Control mice received untreated water. For histological analysis, colons were fixed overnight at 4°C in Bouin's solution (Fisher Scientific 112016), embedded in paraffin by the UT Southwestern Histology Core, and stained with hematoxylin and eosin. Disease severity was assessed as previously described (*46*).

**Flow cytometry**

Lamina propria cells were isolated from mouse colon using a published protocol (*47, 48*). Briefly, colon tissues were cut into small pieces and washed with ice-cold PBS containing 5% fetal bovine serum (PBS-FBS). Epithelial cells were removed by incubating intestinal tissues in Hank's buffered salt solution (HBSS) supplemented with 2 mM EDTA and 1 mM DTT, followed by extensive washing with PBS-FBS. Remaining tissue was washed again in cold PBS and digested with 0.025 mg/ml collagenase, 0.05 mg/ml DNase I, and 0.5 U/ml dispase to generate single-cell suspensions. For flow cytometry, cells were stained with antibodies listed in S5. Samples were analyzed using a NovoCyte Flow Cytometer (ACEA Biosciences), and data were processed with FlowJo software.

#### *Salmonella* Typhimurium infections of mice

*S*. Typhimurium-GFP (gift from Dr. Vanessa Sperandio)(*49*) was grown in Luria broth at 37°C with 50 μg/ml streptomycin and 100 μg/ml ampicillin. Wild-type and *Hdac5*$^{-/-}$ mice or *Hdac5*$^{fl/fl}$ and *Hdac5*$^{\Delta IEC}$ mice were infected by intragastric gavage with 5 × $10^9$ CFU per mouse. After 24 hours, 1 μl of lumen chyme of small intestine and colon were diluted in 1 ml PBS buffer. Small intestine, colon, liver, spleen, and mesenteric lymph nodes (MLN) were homogenized in PBS, and bacterial loads were quantified by dilution plating on Luria broth agar containing the same antibiotics.

#### Chromatin immunoprecipitation (ChIP)

Mouse intestines (ileum or colon) tissues were washed with ice-cold PBS and epithelial cells were recovered as described above. ChIP-qPCR was carried out as previously described (*13*). Briefly, cells were washed twice with ice-cold PBS and fixed in 1% formaldehyde at room temperature for 10 minutes, then quenched with 125 mM glycine for 10 minutes. After two PBS washes, cells were resuspended in 0.5 ml lysis buffer (20 mM Tris-HCl pH 8, 60 mM KCl, 1 mM EDTA, 0.5% NP-40, protease inhibitors) and incubated at 4°C for 15 minutes. Nuclei were pelleted and lysed in 250 μl RIPA buffer (Thermo Scientific 89900) with protease inhibitors, then sonicated (Bioruptor Pico, Diagenode; 20 cycles, 30 sec on/off). The supernatant was pre-cleared with 10 μl protein G magnetic beads for one hour and incubated overnight at 4°C with 3 μg of anti-H3K9ac or anti-H3K27ac antibodies (Table S5). Immunocomplexes were captured with 50 μl protein G beads for 1.5 hours at 4°C, washed four times with LiCl wash buffer (100 mM Tris-HCl pH 7.5, 500 mM LiCl, 0.5% NP-40, 0.5% sodium deoxycholate), and rinsed with TE buffer (10 mM Tris-HCl pH 8, 1 mM EDTA). DNA was eluted in 150 μl TES buffer (TE with 1% SDS, 150 mM NaCl, 5 mM DTT) by incubating beads at 65°C for 8 hours and purified using the ChIP DNA Clean & Concentrator kit (Zymo Research D5205). ChIP DNA was analyzed by qPCR, and enrichment was calculated by normalizing signals to input DNA and to control regions lacking known binding sites near the gene of interest.

**Generation of *HDAC5*$^{-/-}$ HT-29 cells**

Single guide RNAs (sgRNAs) targeting exons 3 and 10 of *Hdac5* were cloned into the LentiCRISPR v2 plasmid (Table S4). Lentiviral particles were generated by transfecting HEK293T cells with the sgRNA-encoding LentiCRISPR v2 construct along with packaging plasmids psPAX2 and pMD2.G using FuGENE® HD Transfection Reagent (Promega, E2311), according to the manufacturer's protocol. Viral supernatants were harvested 48 hours post-transfection, filtered through a 0.45 µm membrane, and used to transduce HT-29 cells in the presence of 8 µg/ml polybrene.

Twenty-four hours after transduction, HT-29 cells were selected with puromycin (2 µg/ml) for 5–7 days. Following selection, single-cell clones were isolated by serial dilution and seeded into 96-well plates. Clonal populations were screened by PCR (primer sequences are listed in Table S3) and verified by Sanger sequencing. Control HT-29 cells were generated using the same protocol with a non-targeting sgRNA. The sgRNA oligonucleotide sequences are provided in Table S3.

**Subcellular fractionation for HDAC5 localization**

To assess the subcellular localization of HDAC5, nuclear and cytoplasmic extracts were prepared using NE-PER™ Nuclear and Cytoplasmic Extraction Reagents (Thermo Scientific, PI78833). For mouse IECs, ileal tissues were washed in ice-cold PBS and IECs were isolated using 10 mM EDTA, as described above. For HT-29 cells, HA-tagged human HDAC5 was cloned into the pMSCV-Blasticidin vector (Table S4) and transfected into wild-type and *HDAC5*$^{-/-}$ HT-29 cells using FuGENE® HD Transfection Reagent (Promega, E2311). Transfected cells were harvested by trypsinization, washed with PBS, and pelleted by centrifugation. For each sample, 20 µl of the cell pellet was used for fractionation. An additional 20 µl was lysed in 100 µl of T-PER (Thermo Scientific, 78510) to generate a whole cell lysate control.

Cytoplasmic and nuclear extraction was carried out according to the manufacturer's instructions. Briefly, 100 µl of ice-cold CER I buffer was added to each 20 µl pellet, and the samples were vortexed vigorously for 15 seconds and incubated on ice for 10 minutes. Then, 11 µl of ice-cold CER II buffer was added, followed by 5 seconds of vortexing and 1 minute on ice. This vortex/incubation cycle was repeated once, then samples were centrifuged at maximum speed for 5 minutes. The supernatant (cytoplasmic extract) was transferred to a pre-chilled tube. The remaining pellet was resuspended in 100 µl of ice-cold NER buffer, vortexed for 15 seconds, and incubated on ice for 10 minutes. This vortex/incubation step was repeated four times, followed by centrifugation at maximum speed for 10 minutes. The supernatant (nuclear extract) was collected in a separate pre-chilled tube. All fractions were mixed with 5X SDS-PAGE loading buffer and analyzed by immunoblotting for HDAC5.

**Rapamycin treatment**

Rapamycin stock solution (10 mg/ml in DMSO; Santa Cruz Biotechnology) was diluted to 1 mg/ml in dimethyl sulfoxide (DMSO; Sigma-Aldrich, D2438) containing 5% PEG400 and 5% Tween-80. Mice were injected intraperitoneally with rapamycin at a dose of 6 mg/kg for 3 consecutive days. For in vitro experiments, LS513 cells were seeded in 6-well plates at a density of $6 \times 10^5$ cells/ml in 2 ml of medium per well. The following day, rapamycin was added at final concentrations of 200 nM or 500 nM by adding 4 µl or 10 µl respectively, of a 100 µM solution prepared in DMSO. DMSO was added to control wells as a vehicle control. Plates were gently mixed and incubated at 37°C for 6 hours.

**Polysome profiling assay for translation efficiency**

IECs were isolated from conventional wild-type, conventional *Hdac5*$^{-/-}$, or germ-free wild-type mice using 10 mM EDTA and 100 µg/ml cycloheximide. Cells were lysed in polysome extraction buffer (20 mM Tris-HCl, pH 7.5, 100 mM KCl, 5 mM $MgCl_2$, 0.5% NP-40, 100 µg/ml cycloheximide, 1 mM DTT) supplemented with protease inhibitors (Roche, 11697498001) and RNase inhibitors (Life Tech, 10777019). Nuclei were pelleted by centrifugation, and the resulting supernatants were layered onto 10–50% sucrose gradients and centrifuged at 190,000 × *g* for 90 minutes. Gradients were fractionated into five equal fractions, and RNA was extracted from each fraction using QIAzol reagent (Qiagen, 79306). Ribosome-associated mRNAs were quantified by qPCR, and transcript levels of barrier defense genes were normalized to *Gapdh*.

**Co-immunoprecipitation assays**

Co-immunoprecipitation assays were performed using lysates from either small intestinal epithelial cells or HT-29 cells. For HT-29 cells, HA-tagged HDAC5 (cloned into pMSCV-Blasticidin; Table S4) or HA-tagged 14-3-3 (Table S4) was transfected using FuGene® HD Transfection Reagent (Promega, E2311). Forty-eight hours post-transfection, cells were collected and lysed in T-PER buffer (Thermo Scientific, 78510). Protein concentrations were determined using a NanoDrop One spectrophotometer (Thermo Scientific) and adjusted to 5 mg/ml. Lysates were incubated with 2 µl anti-HA antibody (Cell Signaling Technology, 3724S) at 4°C for at least 1 hour, followed by incubation with 20 µl Pierce™ Protein A Agarose beads (Thermo Scientific, 20333) overnight at 4°C. Beads were washed three times with T-PER buffer, then resuspended in 50 µl of SDS-PAGE loading buffer and boiled for 5 minutes prior to SDS-PAGE separation. Immunoblotting or silver staining was used to analyze proteins in the immunoprecipitation pellets.

For coimmunoprecipitations using small intestinal epithelial cell lysates, lysates were prepared as described in the immunoblotting section, and antibodies used for immunoprecipitation are listed in Table S5.

**Liquid chromatography–tandem mass spectrometry (LC–MS/MS)**

Proteins enriched in immunoprecipitation pellets were separated by SDS-PAGE. Protein bands were silver stained with Pierce™ Silver Stain for Mass Spectrometry (Thermo Scientific #24600). The indicated bands were cut into 1 $mm^3$ cubes and analyzed by LC–MS/MS by the UT Southwestern Proteomics Core. Samples were digested overnight with trypsin (Pierce) following reduction and alkylation with DTT and iodoacetamide (Sigma–Aldrich). The samples then underwent solid-phase extraction cleanup with an Oasis HLB plate (Waters) and were subsequently dried and reconstituted into 10 µl of 2% ACN, 0.1% TFA. 5 µl of these samples was injected onto a QExactive HF mass spectrometer coupled to an Ultimate 3000 RSLC-Nano liquid chromatography system. Samples were injected onto a 75 µm inner diameter, 15-cm long EasySpray column (Thermo Scientific) and eluted with a gradient from 0-28% buffer B over 90 min with a flow rate of 250 nl/min. Buffer A contained 2% (v/v) ACN and 0.1% formic acid in water, and buffer B contained 80% (v/v) ACN, 10% (v/v) trifluoroethanol, and 0.1% formic acid in water. The mass spectrometer operated in positive ion mode with a source voltage of 2.4 kV and an ion transfer tube temperature of 275°C. Mass spectrometry scans were acquired at 120,000 resolution in the Orbitrap and up to 20 MS/MS spectra were obtained for each full spectrum acquired using higher-energy collisional dissociation (HCD) for ions with charges 2-8. Dynamic exclusion was set for 20 seconds after an ion was selected for fragmentation. Raw MS data files were analyzed using Proteome Discoverer v2.4 (Thermo Scientific), with peptide identification performed using Sequest HT searching against the human protein database from UniProt (downloaded on March 12, 2020). Fragment and precursor tolerances of 10 ppm and 0.02 Da were specified, and three missed cleavages were allowed. Carbamidomethylation of Cys was set as a fixed modification, with oxidation of Met and acetylation of Lys set as a variable modification. The false-discovery rate (FDR) cutoff was 1% for all peptides.

**Intestinal organoid culture**

Organoids were generated from small intestinal crypts isolated from 6- to 8-week-old mice, using a protocol adapted from Sato et al. (*50*). Small intestines were rinsed with PBS, fat was removed, and the tissue was opened longitudinally and cut into 5–10 cm segments. These segments were washed in PBS, and villi were removed by firmly scraping the luminal surface with a glass microscope slide. The intestine was then cut into ~1 cm pieces and transferred to 20 ml cold PBS, followed by vigorous shaking. This wash step was repeated with fresh PBS until the supernatant was clear.

Tissue pieces were then incubated in 20 ml of 5 mM EDTA in PBS on a rotator at 4°C for 15 minutes to loosen crypts. Crypts were released by vigorous shaking, and successful elution was confirmed by microscopy. Supernatants were pooled and centrifuged at 300 × *g* for 3 minutes. Pellets were resuspended in ADF medium (DMEM + GlutaMAX™ [Gibco]) supplemented with 10% FBS and passed through a 70 µm cell strainer. Centrifugation and resuspension were repeated 2–3 times until the supernatant was clear.

Crypts were counted using trypan blue exclusion and a light microscope. The final pellet was resuspended in 50 µl IntestiCult™ Organoid Growth Medium (StemCell Technologies, 06005) and mixed with 200 µl Matrigel (BD Biosciences, 356231). Aliquots of the mixture were plated into six wells of a pre-warmed 24-well plate, typically yielding 100–500 organoids per well. Plates were incubated at 37°C for 5–10 minutes to allow Matrigel polymerization, followed by the addition of 500 µl IntestiCult™ medium per well. Organoids were maintained at 37°C with 5% $CO_2$, and culture medium was refreshed every 2–3 days.

To prepare luminal contents, a ~5 cm segment of ileum was flushed with PBS. The flushed luminal contents were collected, centrifuged to remove debris, and the supernatant was passed through a 0.45 µm syringe filter (Thermo Scientific). The resulting filtrate was used immediately for organoid treatment.

Organoids were cultured for 7–10 days. To harvest them, 500 µl of cold medium (DMEM + 10% FBS) was added to each well, and the contents were transferred to Eppendorf tubes after Matrigel dissolution. Organoids were collected by centrifugation and mechanically dissociated by pipetting with bent 200 µl tips. The disrupted organoids were washed with fresh medium and replated into wells with 200 µl of IntestiCult™ Organoid Growth Medium (StemCell Technologies). For treatment, 10 µl of filtered ileal lumen supernatant was added to each well. Control wells received 10 µl of PBS. Organoids were incubated for 4 hours, then pelleted by centrifugation at 300 × *g*. Both supernatants and organoid pellets were collected and analyzed for barrier defense protein levels by immunoblotting.

**Statistical analysis**

Statistical methods and definitions of significance are described in the figure legends. Unless otherwise noted (Fig. 2L; fig. S2C), data are presented as mean ± SEM. Sample sizes were not predetermined using statistical methods. While formal randomization procedures were not employed, mice were assigned to experimental groups at random and samples were processed in no predetermined order. All statistical analyses were carried out using GraphPad Prism. Comparisons between two groups of mice were made using two-tailed Student's *t* tests, with P values less than 0.05 considered statistically significant. Mice that died during the study were the only animals excluded from analysis.

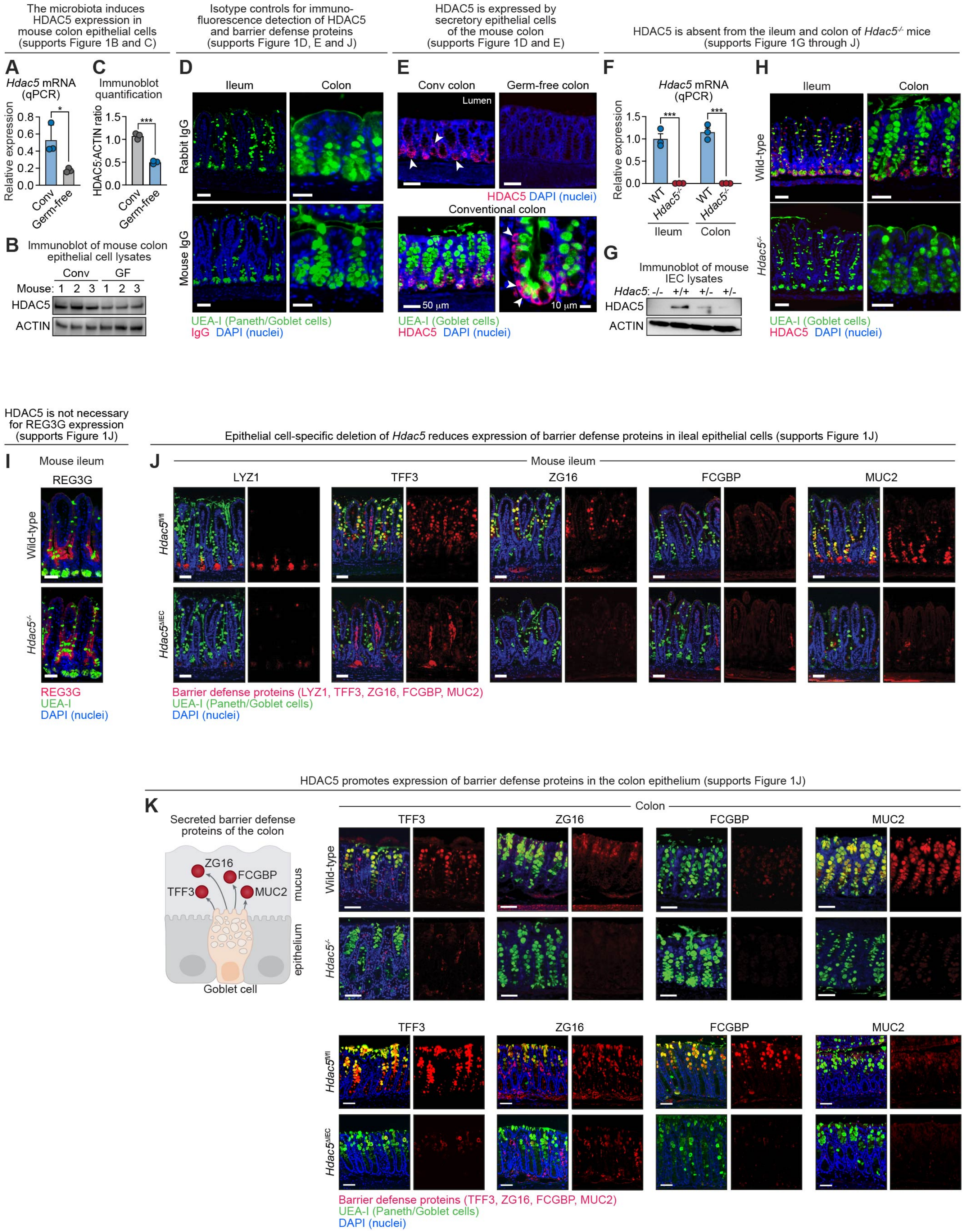

The microbiota induces HDAC5 expression in mouse colon epithelial cells (supports Figure 1B and C)
Isotype controls for immuno-fluorescence detection of HDAC5 and barrier defense proteins (supports Figure 1D, E and J)
HDAC5 is expressed by secretory epithelial cells of the mouse colon (supports Figure 1D and E)
HDAC5 is absent from the ileum and colon of Hdac5-/- mice (supports Figure 1G through J)
A
Hdac5 mRNA (qPCR)
Relative expression
0.0 0.2 0.4 0.6 0.8
*
Conv
Germ-free
C
Immunoblot quantification
HDAC5:ACTIN ratio
0.0 0.5 1.0 1.5
***
Conv
Germ-free
B
Immunoblot of mouse colon epithelial cell lysates
Conv GF
Mouse: 1 2 3 1 2 3
HDAC5
ACTIN
D
Ileum
Colon
Rabbit IgG
Mouse IgG
UEA-I (Paneth/Goblet cells)
IgG DAPI (nuclei)
E
Conv colon
Germ-free colon
Lumen
HDAC5 DAPI (nuclei)
Conventional colon
50 μm
10 μm
UEA-I (Goblet cells)
HDAC5 DAPI (nuclei)
F
Hdac5 mRNA (qPCR)
Relative expression
0.0 0.5 1.0 1.5
***
***
WT
Hdac5-/-
WT
Hdac5-/-
Ileum
Colon
G
Immunoblot of mouse IEC lysates
Hdac5: -/- +/+ +/- +/-
HDAC5
ACTIN
H
Ileum
Colon
Wild-type
Hdac5-/-
UEA-I (Goblet cells)
HDAC5 DAPI (nuclei)
HDAC5 is not necessary for REG3G expression (supports Figure 1J)
I
Mouse ileum
REG3G
Wild-type
Hdac5-/-
REG3G
UEA-I
DAPI (nuclei)
Epithelial cell-specific deletion of Hdac5 reduces expression of barrier defense proteins in ileal epithelial cells (supports Figure 1J)
J
Mouse ileum
LYZ1
TFF3
ZG16
FCGBP
MUC2
Hdac5fl/fl
Hdac5ΔIEC
Barrier defense proteins (LYZ1, TFF3, ZG16, FCGBP, MUC2)
UEA-I (Paneth/Goblet cells)
DAPI (nuclei)
HDAC5 promotes expression of barrier defense proteins in the colon epithelium (supports Figure 1J)
K
Secreted barrier defense proteins of the colon
ZG16
FCGBP
TFF3
MUC2
mucus
epithelium
Goblet cell
Colon
TFF3
ZG16
FCGBP
MUC2
Wild-type
Hdac5-/-
TFF3
ZG16
FCGBP
MUC2
Hdac5fl/fl
Hdac5ΔIEC
Barrier defense proteins (TFF3, ZG16, FCGBP, MUC2)
UEA-I (Paneth/Goblet cells)
DAPI (nuclei)

**Figure S1. HDAC5 promotes expression of barrier defense proteins in intestinal secretory epithelial cells (supports Figure 1).**

**(A)** qPCR of *Hdac5* transcripts in colon epithelial cells of conventional and germ-free mice.
**(B)** Immunoblot detection of HDAC5 in colon epithelial cell lysates from conventional and germ-free mice. ACTIN was the loading control.
**(C)** Band intensities in (B) were measured by scanning densitometry and protein ratios were calculated.
**(D)** Isotype controls for immunofluorescence microscopy in mouse intestine. Scale bars, 50 µm.
**(E)** Upper panels show immunofluorescence microscopy of HDAC5 in colon epithelial cells of conventional and germ-free mice. Lower panels show that HDAC5 is expressed by secretory epithelial cells (goblet cells) of the mouse colon. Scale bars, 50 µm.
**(F)** qPCR analysis of *Hdac5* transcripts in ileum and colon epithelial cells of wild-type and *Hdac5*$^{-/-}$ mice.
**(G)** Immunoblot detection of HDAC5 in small intestine IEC lysates from *Hdac5* wild-type (+/+), heterozygotes (+/-) and homozygotes (-/-). ACTIN was the loading control.
**(H)** Immunofluorescence microscopy of HDAC5 in the ileum and colon of wild-type and *Hdac5*$^{-/-}$ mice. Scale bars, 50 µm.
**(I)** Immunofluorescence microscopy of REG3G in the ileum of wild-type and *Hdac5*$^{-/-}$ mice. Scale bars, 50 µm.
**(J)** Immunofluorescence microscopy of barrier defense proteins (LYZ1, TFF3, ZG16, FCGBP, and MUC2) in small intestines of mice with an epithelial cell-specific deletion of *Hdac5* (*Hdac5*$^{\Delta IEC}$ compared to *Hdac5*$^{fl/fl}$ littermate controls). Scale bars, 50 µm.
**(K)** Immunofluorescence microscopy of barrier defense proteins (TFF3, ZG16, FCGBP, and MUC2) in the colons of wild-type and *Hdac5*$^{-/-}$ mice, and *Hdac5*$^{fl/fl}$ and *Hdac5*$^{\Delta IEC}$ mice. Scale bars, 50 µm.

Conv, conventional; GF, germ-free; IEC, intestinal epithelial cells; UEA-I, *Ulex europaeus* agglutinin I; LYZ1, lysozyme; ZG16, zymogen granule 16; TFF3, trefoil factor 3; FCGBP, Fc gamma binding protein; MUC2, mucin 2. Each experiment was performed at least twice; each bar graph data point represents one mouse; each immunoblot lane represents one mouse. Means ± SEM are plotted; *p < 0.05; ***p < 0.001; ns, not significant by Student's *t* test.

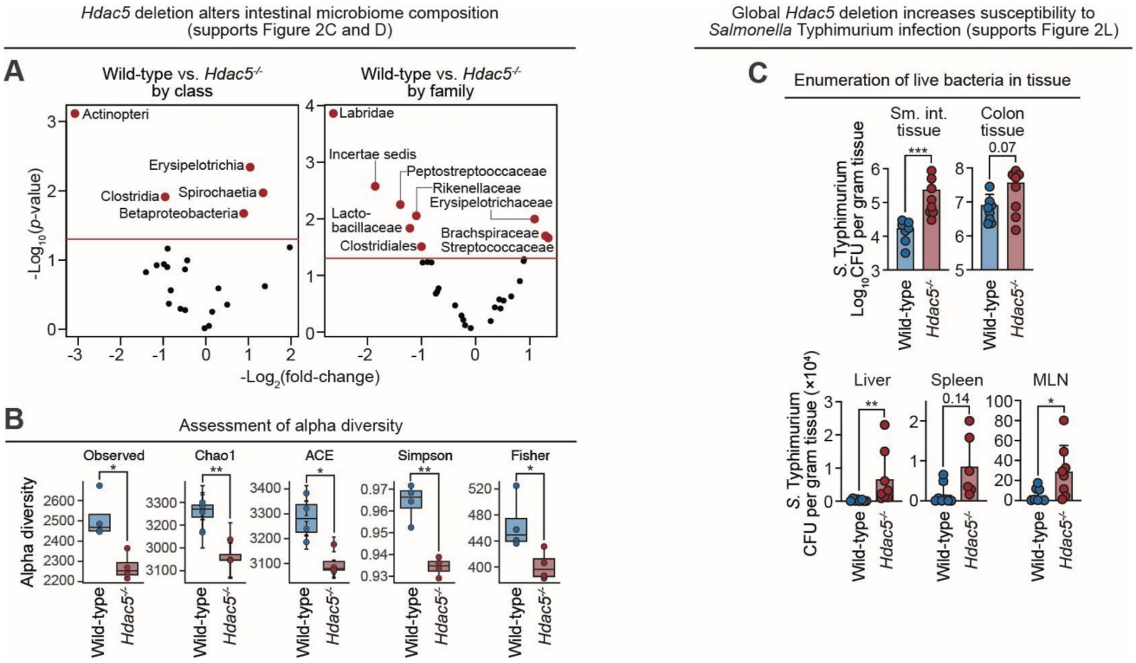

**Figure S2. *Hdac5* deletion alters gut microbiome composition and increases susceptibility to *Salmonella* Typhimurium infection (supports Figure 2).**

**(A)** Volcano plot showing bacterial classes or families with differential abundances in wild-type and *Hdac5*$^{-/-}$ mice. N=4 mice per group.

**(B)** Alpha diversity measurement in fecal bacterial populations wild-type and *Hdac5*$^{-/-}$ mice. The mice were littermates of heterozygous crosses that remained cohoused. Each dot represents one mouse. Means ± SEM are plotted.

**(C)** Global deletion of *Hdac5* increases susceptibility of mice to *Salmonella* Typhimurium infection. Mice were orally inoculated with $5 \times 10^9$ CFU of *Salmonella* Typhimurium. Tissues were collected after 24 hours and analyzed by dilution plating to determine *S*. Typhimurium burdens in the tissues of wild-type and *Hdac5*$^{-/-}$ littermates. N=8 mice/group. CFU, colony forming units; MLN, mesenteric lymph nodes. Geometric means ± SEM are plotted. *p < 0.05; **p < 0.01; ***p < 0.005; ns, not significant by Student's *t* test.

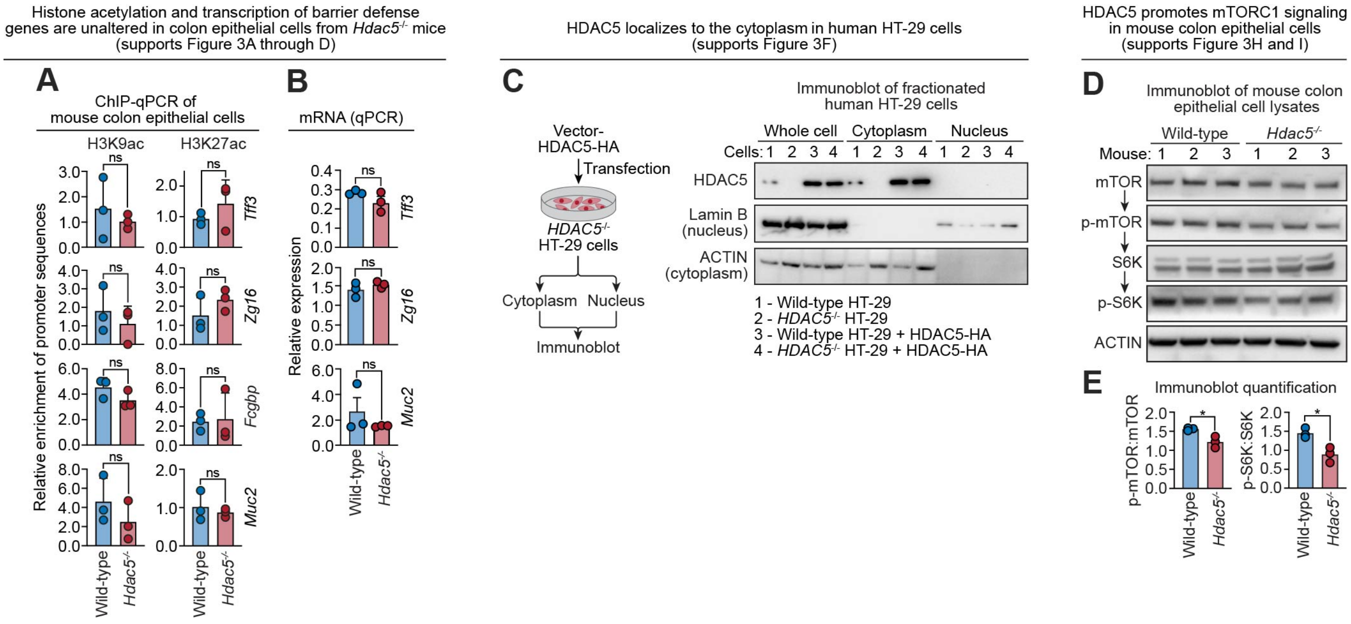

**Figure S3. Histone acetylation and transcription of barrier defense genes are unaltered in colon epithelial cells from *Hdac5*$^{-/-}$ mice (supports Figure 3).**

**(A)** ChIP-qPCR analysis of H3K9ac and H3K27ac at the promoters of *Muc2*, *Tff3*, *Zg16* and *Fcgbp* in colon epithelial cells from wild-type and *Hdac5*$^{-/-}$ mice. N=3 per group. Each data point represents one mouse.

**(B)** qPCR of barrier defense gene transcripts in colon epithelial cells from wild-type and *Hdac5*$^{-/-}$ mice. N=3 mice/group. Each data point represents one mouse.

**(C)** Endogenous or ectopically expressed HDAC5 localizes to the cytoplasm in human colorectal carcinoma HT-29 cells. HT-29 cells were grown in culture and separated by centrifugation into cytoplasm- and nucleus-containing fractions. The fractions were immunoblotted and HDAC5 was detected alongside Lamin B (a marker for nuclei) and ACTIN (a marker for cytoplasm). Lane 1: Wild-type HT-29 cells; Lane 2: *HDAC5*$^{-/-}$ HT-29 cells; Lane 3: wild-type HT-29 cells expressing HDAC5-HA; Lane 4: *HDAC5*$^{-/-}$ HT-29 cells expressing HDAC5-HA.

**(D)** Immunoblot detection of mTORC1 pathway proteins in colon epithelial cell lysates from wild-type and *Hdac5*$^{-/-}$ mice. Each lane represents one mouse.

**(E)** Band intensities from (D) were measured by scanning densitometry and protein ratios were calculated.

Each experiment was performed at least twice; means ± SEM are plotted; *p < 0.05; ns, not significant by Student's *t* test.

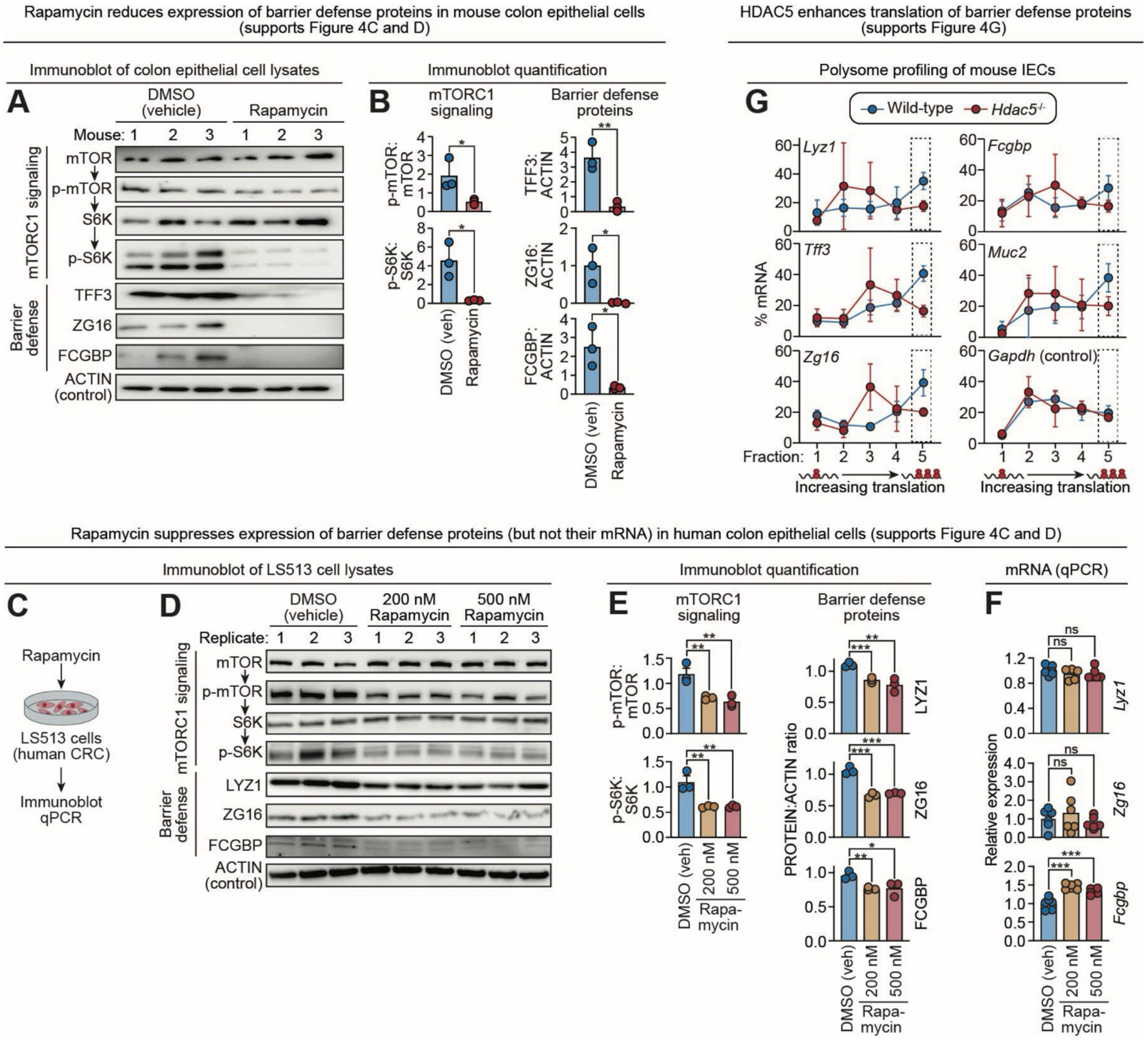

**Figure S4. HDAC5 promotes barrier defense protein translation through mTOR (supports Figure 4).**

**(A)** Immunoblot of colon epithelial cell lysates from mice treated with rapamycin or DMSO (vehicle control) with detection of mTORC1 signaling pathway components and barrier defense proteins. ACTIN was the loading control.

**(B)** Band intensities from (A) were measured by scanning densitometry and protein ratios were calculated.

**(C)** LS513 cells (a human colorectal carcinoma cell line) were treated with Rapamycin at 200 nM and 500 nM for 6 hours. DMSO was the vehicle control.

**(D)** LS513 cells treated with rapamycin or DMSO were collected, lysed, and blotted. Blots were detected with antibodies against mTORC1 signaling pathway proteins and barrier defense proteins. ACTIN was the loading control. Each lane represents an independent replicate.

**(E)** Immunoblot band intensities from (C) were measured by scanning densitometry and protein ratios were calculated.

**(F)** qPCR of barrier defense gene transcripts. N=6 independent replicates per group.

**(G)** Polysome profiling of barrier defense gene transcripts in small intestine epithelial cells from wild-type and *Hdac5*$^{-/-}$ mice. The percentage of transcripts from each gradient fraction are shown by line graphs. Fraction 5 contains mRNA associated with the highest number of ribosomes, indicating a high rate of translation. *Gapdh* transcripts were analyzed as a control. Bar graphs representing Fraction 5 from this experiment are shown in Figure 4G.

LYZ1, lysozyme; TFF3, trefoil factor 3; ZG16, zymogen granule 16; FCGBP, Fc gamma binding protein. Each experiment was performed at least twice; each bar graph data point represents one mouse or one cell culture replicate; each line graph data point represents the mean from three mice; each immunoblot lane represents one mouse or one cell culture replicate. Means ± SEM are plotted; $*p < 0.05$; $**p < 0.01$; $***p < 0.001$; ns, not significant by Student's *t* test.

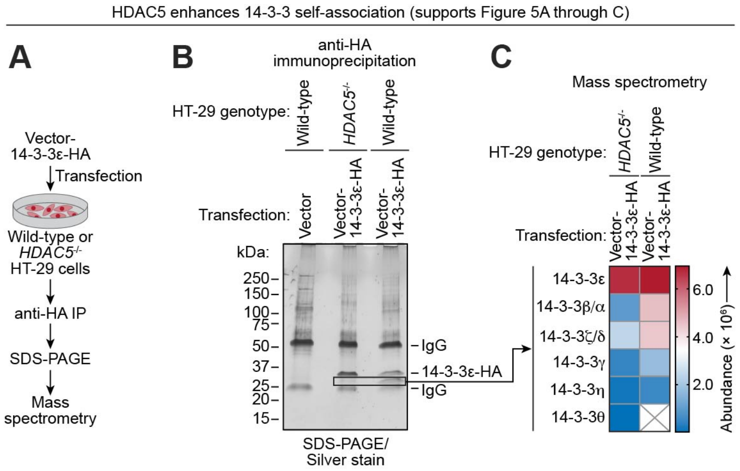

**Figure S5. HDAC5 enhances 14-3-3 self-association (supports Figure 5).**

**(A)** To test for 14-3-3 self-association, we used HA-tagged 14-3-3ε so that we could distinguish it from endogenous 14-3-3. A plasmid encoding HA-tagged 14-3-3ε was transfected into HT-29 cells (wild-type or *HDAC5*$^{-/-}$), and cell lysates were subjected to immunoprecipitations with anti-HA antibody. Proteins in the anti-HA immunoprecipitation pellets were separated by SDS-PAGE and protein bands were visualized by silver staining. Protein bands enriched in anti-HA immunoprecipitation pellets were analyzed by LC–MS/MS.

**(B)** Silver stained SDS-PAGE of immunoprecipitation pellets. The rectangle highlights the protein bands that were analyzed by LC–MS/MS.

**(C)** Heat map of proteins identified by mass spectrometry as being associated with HA-tagged 14-3-3ε. The list is ordered by protein abundance. The results indicate that HA-tagged 14-3-3ε associates with more endogenous 14-3-3 proteins in wild-type HT-29 cells, suggesting that self-association decreases when HDAC5 is absent.

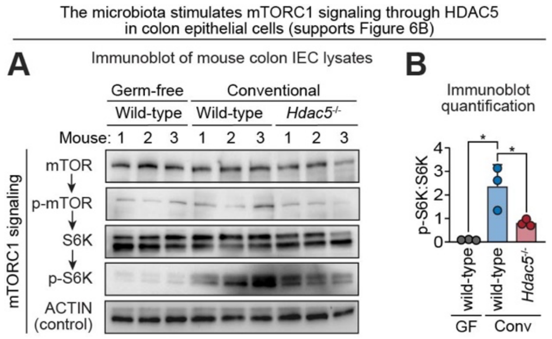

**Figure S6. The microbiota stimulates mTORC1 signaling through HDAC5 in colon epithelial cells (supports Figure 6).**

**(A)** Immunoblot of colon epithelial cell lysates from conventional mice (wild-type, *Hdac5*$^{-/-}$) and germ free (wild-type) mice, with antibody detection of mTORC1 signaling pathway components. Each data point represents one mouse.

**(B)** Band intensities in (A) were measured by scanning densitometry and protein ratios were calculated. Each data point represents one mouse. Means ± SEM are plotted; *p<0.05 by Student's *t* test.

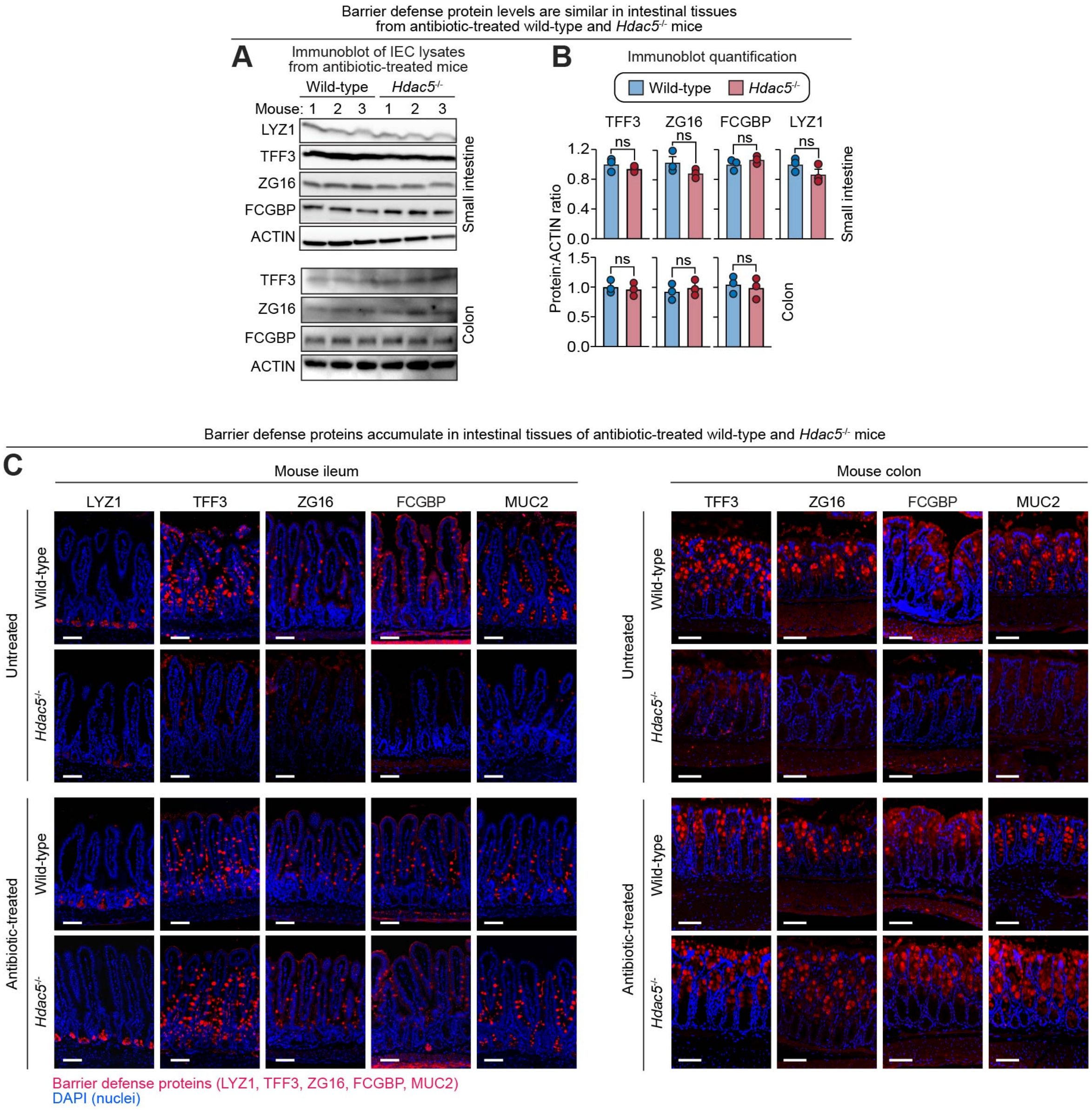

**Figure S7. Intestinal barrier defense protein levels appear similar in antibiotic-treated wild-type and *Hdac5*$^{-/-}$ mice.**

**(A)** Immunoblot of small intestine and colon epithelial cell lysates from antibiotic-treated mice (wild-type, *Hdac5*$^{-/-}$), with detection of barrier defense proteins. ACTIN was the loading control. Each lane represents one mouse.

**(B)** Band intensities were measured by scanning densitometry and protein ratios were calculated. Each data point represents one mouse. Means ± SEM are plotted; ns, not significant by Student's *t* test.

**(C)** Immunofluorescence microscopy of barrier defense proteins in the small intestines and colons of wild-type and *Hdac5*$^{-/-}$ mice. Mice were either treated with antibiotics or left untreated. Scale bars, 50 µm.

IEC, intestinal epithelial cells; LYZ1, lysozyme; ZG16, zymogen granule 16; TFF3, trefoil factor 3; FCGBP, Fc gamma binding protein; MUC2, mucin 2.

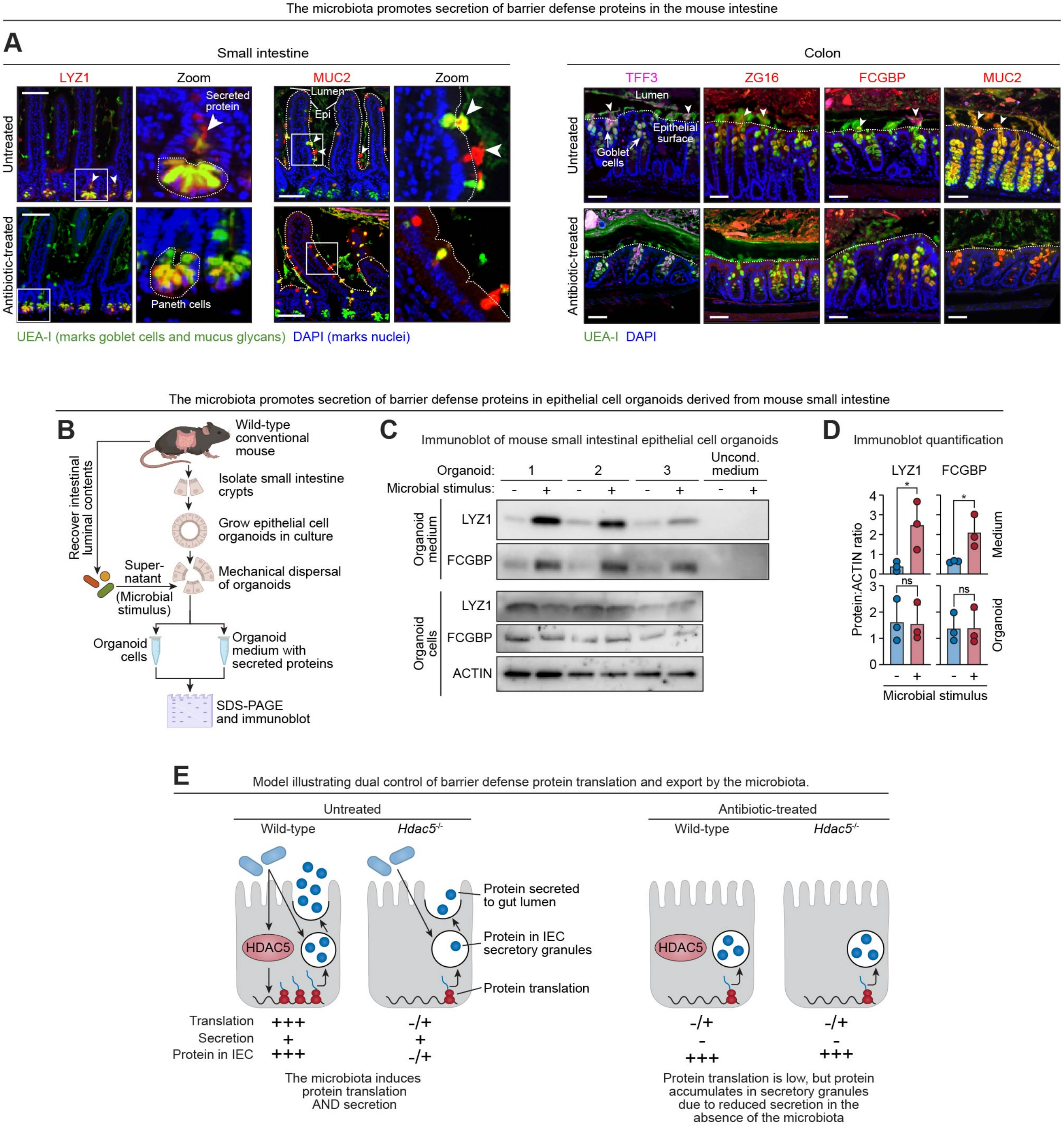

**Figure S8. The microbiota regulates both translation and secretion of barrier defense proteins.**

**(A)** Mouse small intestines or colons, with luminal contents intact, were fixed in methacarn solution to preserve the mucus layer. Barrier defense proteins were visualized by immunofluorescence microscopy to assess their secretion into the mucus. Arrowheads indicate secreted proteins localized within the mucus layer. Dashed white lines outline the edges of intestinal epithelial cells. Scale bars, 50 µm.

**(B)** Workflow showing the use of organoid cultures to quantify the secretion of barrier defense proteins in response to a microbial stimulus. Epithelial cell organoids were established from small intestinal crypt cells recovered from wild-type mice. The organoids were disrupted to expose the luminal (apical) surfaces of the organoid epithelial cells. Supernatants from intestinal luminal contents recovered from conventional mice were added to the culture (PBS was used as a control). After two hours, organoids and culture supernatants were collected and analyzed by immunoblot for secretion of barrier defense proteins.

**(C)** Immunoblots of organoid cells or culture supernatants were detected with antibodies directed against FCGBP and LYZ. ACTIN was the loading control. Each organoid was derived from an individual mouse. For the unconditioned (uncond.)

medium control, luminal contents were added to fresh culture medium that was not associated with organoids. This was to control for whether the immunoblot bands arose from the luminal contents.

**(D)** Band intensities in (C) were measured by scanning densitometry and protein ratios were calculated. Means ± SEM are plotted; *p<0.05; ns not significant by Student's *t* test.

**(E)** ***Model illustrating dual control of barrier defense protein translation and export by the microbiota.*** Our finding that the microbiota enhances translation of mucus barrier proteins presents a paradox alongside the observation of similar tissue levels of these proteins in conventional and antibiotic-treated mice (Fig. S7 and (*10*)). The paradox is explained by simultaneous control of protein translation and export by the microbiota. Thus, steady-state epithelial protein levels reflect the net effect of production and export. In untreated wild-type mice, the microbiota induces *Hdac5* expression, which enhances barrier defense protein translation via mTORC1 signaling. Simultaneously, the microbiota promotes secretion of these proteins. In *Hdac5*$^{-/-}$ mice with an intact microbiota, translation is reduced but secretion remains active, resulting in low intracellular protein levels. In contrast, antibiotic-treated mice (regardless of genotype) exhibit low *Hdac5* expression and reduced translation, but also impaired secretion. As a result, although the proteins are translated at a low rate, they accumulate within epithelial cells even in the absence of the microbiota. This likely enables rapid deployment of these proteins in response to a microbial signal. Dual control of protein synthesis and export by the microbiota thus ensures that barrier defense protein production is matched to demand at the barrier surface.

IEC, intestinal epithelial cells; LYZ1, lysozyme; ZG16, zymogen granule 16; TFF3, trefoil factor 3; FCGBP, Fc gamma binding protein; MUC2, mucin 2.

**Table S1. Liquid chromatography–tandem mass spectrometry analysis of HDAC5-interacting proteins (supports Figure 5C)**

| Accession | Description | #PSMs[1] | MW (kDa) | Abundance in *HDAC5*$^{-/-}$ cells | Abundance in *HDAC5*$^{-/-}$ cells + HDAC5-HA | Fold enrichment |
|---|---|---|---|---|---|---|
| P62258 | 14-3-3 protein epsilon | 344 | 29.2 | 8.25E+06 | 7.21E+07 | 8.7 |
| P63104 | 14-3-3 protein zeta/delta | 94 | 27.7 | 1.97E+06 | 1.71E+07 | 8.7 |
| Q04917 | 14-3-3 protein eta | 87 | 28.2 | 1.38E+06 | 1.15E+07 | 8.3 |
| P31946 | 14-3-3 protein beta/alpha | 97 | 28.1 | 7.51E+05 | 4.51E+06 | 6.0 |
| P35527 | Keratin, type I cytoskeletal 9 | 59 | 62.0 | 7.47E+05 | 4.46E+06 | 6.0 |
| P61981 | 14-3-3 protein gamma | 95 | 28.3 | 1.29E+06 | 7.63E+06 | 5.9 |
| P27348 | 14-3-3 protein theta | 52 | 27.7 | 2.22E+05 | 1.29E+06 | 5.8 |
| K7EJL6 | Histone deacetylase 5 (Fragment) | 17 | 8.3 | 1.86E+05 | 7.54E+05 | 4.1 |
| Q02978 | Mitochondrial 2-oxoglutarate/malate carrier protein | 15 | 34.0 | 2.49E+05 | 7.73E+05 | 3.1 |
| P13647 | Keratin, type II cytoskeletal 5 | 25 | 62.3 | 1.82E+05 | 5.44E+05 | 3.0 |
| P62701 | 40S ribosomal protein S4, X isoform | 39 | 29.6 | 2.81E+06 | 7.43E+06 | 2.6 |
| P04264 | Keratin, type II cytoskeletal 1 | 138 | 66.0 | 3.33E+06 | 8.58E+06 | 2.6 |
| P62241 | 40S ribosomal protein S8 | 18 | 24.2 | 8.42E+05 | 2.07E+06 | 2.5 |
| P02538 | Keratin, type II cytoskeletal 6A | 24 | 60.0 | 2.28E+04 | 4.97E+04 | 2.2 |

[1]PSM, peptide spectrum match

**Table S2. Liquid chromatography–tandem mass spectrometry analysis of 14-3-3 interacting proteins (supports Figure S5C)**

| Acces-sion | Description | Coverage (%) | # Peptides | # PSMs[1] | # Unique peptides | MW (kDa) | Gene symbol | Abundance Wild-type | Abundance *HDAC5*$^{-/-}$ |
|---|---|---|---|---|---|---|---|---|---|
| P62258 | 14-3-3 protein epsilon | 85 | 28 | 264 | 26 | 29.2 | *YWHAE* | 5.31E+06 | 9.09E+06 |
| P63104 | 14-3-3 protein zeta/delta | 60 | 14 | 61 | 9 | 27.7 | *YWHAZ* | 4.54E+06 | 3.62E+06 |
| P31946 | 14-3-3 protein beta/alpha | 66 | 14 | 50 | 7 | 28.1 | *YWHAB* | 4.56E+06 | 1.38E+06 |
| P61981 | 14-3-3 protein gamma | 42 | 10 | 42 | 5 | 28.3 | *YWHAG* | 2.72E+06 | 1.05E+06 |
| Q04917 | 14-3-3 protein eta | 31 | 8 | 36 | 4 | 28.2 | *YWHAH* | 5.53E+05 | 2.08E+05 |
| P27348 | 14-3-3 protein theta | 28 | 6 | 26 | 2 | 27.7 | *YWHAQ* | 6.66E+04 | 2.18E+05 |

[1]PSM, peptide spectrum match

**Table S3. Oligonucleotide sequences used in this study**

| | | |
|---|---|---|
| **Primer sequences for qPCR** | | |
| **Target** | **Forward** | **Reverse** |
| *Hdac5* | GACTTTCCCCTCCGTAAAACG | TGCCATCCTTTCGACGCAG |
| *Lyz1* | GAGACCGAAGCACCGACTATG | CGGTTTTGACATTGTGTTCGC |
| *Muc2* | GACTGACGACAAGAAGAACGTG | AGAGTCACAAAAAGCTGCATGA |
| *Fcgbp* | GGCCCAACTTGGAGAATGGAG | AAGCCAGCATCACAGACACAG |
| *Tff3* | TTGCTGGGTCCTCTGGGATAG | TACACTGCTCCGATGTGACAG |
| *Zg16* | CTCGGCCTCTGCTAATTCCAT | GCACCTGGAGACCTACTATGT |
| *Gapdh* | TGGCAAAGTGGAGATTGTTGCC | AAGATGGTGATGGGCTTCCCG |
| **Primer sequences for ChIP-qPCR** | | |
| **Target** | **Forward** | **Reverse** |
| *Lyz1*_ChIP_pos | AAAGCATGCCTGCTATTTAAGC | CTGGGCTGCTAGAACAACTCTT |
| *Lyz1*_ChIP_neg | CATTTTGTGGCTATGGTTCGTA | TCCCTTTGATGCTTTCTGTTTT |
| *Muc2*_ChIP_pos | TGCTTGGACTCCACATTATCAC | CTCCCTATGAGAGACCCATCTG |
| *Muc2*_ChIP_neg | TGACTCCCACTGTAGCAGAGAA | CCATTGTTTGGATTCCCTATGT |
| *Fcgbp*_ChIP_pos | GGCACTGAACGAAGTGTGTTAG | AAGGATACGAAGGGGAGAAGAC |
| *Fcgbp*_ChIP_neg | ATAGGGAGAGGCCCAAAAGTAG | TTCACTAATAACCCCGGAAATG |
| *Tff3*_ChIP_pos | GCTCAGTGTGTGCTCAGAAATC | AATGGAGGCTCAGAGAGAGTTG |
| *Tff3*_ChIP_neg | CTCTGCTGCTATGAGGGCTATT | CTCACACTCCCCAGTCCTTATC |
| *Zg16*_ChIP_pos | AGCCAAACTGTCCATATCCATC | TGCAACTTCTAGGGGAAAGGTA |
| *Zg16*_ChIP_neg | TTTGTTATTGGGGCACATCATA | TCCGCATAGGAAGAGTAAGAGC |
| **Primers for HA-tagged HDAC5 construct** | | |
| *HDAC5*-F-BglII | GTCAGATCTATGTAC CCA TAC GAT GTT CCA GAT TAC GCT AACTCTCCCAACGAGTCGGC | |
| *HDAC5*-R-XhoI | CAGCTCGAGTCACAGGGCAGGCTCCTGCTC | |
| **Sequences of sgRNAs used to generate *HDAC5*$^{-/-}$ HT-29 cells** | | |
| *HDAC5* exon3_sgRNA | GAATCCTGCGTGGTAGCCTTTGG | |
| *HDAC5* exon10_sgRNA | GATGCTAGTCCAAAGTTGGCAGG | |
| **Genotyping primers for *HDAC5*$^{-/-}$ HT-29 cells** | | |
| **Target** | **Forward** | **Reverse** |
| *HDAC5* validate1 | GTCATGCTGTTTCTGGAACTCA | TTGCTTGGACTTAATTGCATTG |
| *HDAC5* validate2 | TTCTGCTAAGGGGAACTGTGAC | AAGTTCATGGCTTCAGCCTC |
| *HDAC5* validate4 | GAACTCTGGTCCAAAGAAGCAT | GGAACAAGGAGAAGAGCAAAGA |
| **Probe sequences used for 16*S* rRNA fluorescence in situ hybridization (FISH)** | | |
| 16*S* rRNA FISH | [AminoC6+Alexa488]- GCTGCCTCCCGTAGGAGT-[AmC7~Q+Alexa488] | |
| Non-specific FISH probe | [AminoC6+Alexa488]- ACTCCTACGGGAGGCAGC-[AmC7~Q+Alexa488] | |
| **Primers for 16*S* gene sequencing** | | |
| 16*S* V3 – forward primer | TCGTCGGCAGCGTCAGATGTGTATAAGAGACAGCCTACGGGNGGCWGCAG | |
| 16*S* V4 – reverse primer | GTCTCGTGGGCTCGGAGATGTGTATAAGAGACAGGACTACHVGGGTATCT AATCC | |

**Table S4. Plasmids used in this study**

| Plasmid name | Purpose | Source |
|---|---|---|
| pMSCV-Blasticidin | Expression of HA-tagged HDAC5 | Addgene #75085 |
| pcDNA3-HA-14-3-3ε | Expression of HA-tagged 14-3-3ε | Addgene #13273 |
| LentiCRISPR v2 | Generation of *HDAC5*$^{-/-}$ HT-29 cells | Addgene #52961 |
| pMD2.G | Packaging plasmid; generation of *Hdac5*$^{-/-}$ HT-29 cells | Addgene #12259 |
| psPAX2 | Packaging plasmid; generation of *Hdac5*$^{-/-}$ HT-29 cells | Addgene #12260 |

**Table S5. Antibodies used in this study**

| Target protein | Application | Source |
|---|---|---|
| HDAC5 | Immunofluorescence | Active motif #40970 |
| HDAC5 | Immunoblot, immunoprecipitation | Novus #NBP2-22152 |
| MUC2 | Immunofluorescence, immunoblot | Santa Cruz #SC-15334 |
| LYZ1 | Immunofluorescence, immunoblot | Dako #A0099 |
| FCGBP | Immunofluorescence, immunoblot | Novus #NBP1-90462 |
| ZG16 | Immunofluorescence, immunoblot | Proteintech #17397-1-AP |
| TFF3 | Immunofluorescence, immunoblot | Thermo Scientific #14-4758-82 |
| REG3G | Immunofluorescence | Cash et al. (*6*) |
| GFP | Immunofluorescence | Abcam #ab5450 |
| Actin | Immunoblot | Cell Signaling #5125S |
| HA | Immunoprecipitation | Cell Signaling #3724S |
| Lamin B | Immunoblot | Abcam, #ab133741 |
| mTOR | Immunoblot | Cell Signaling #2983S |
| p-mTOR | Immunoblot | Cell Signaling #5536S |
| S6K | Immunoblot | Cell Signaling #9202S |
| p-S6K | Immunoblot | Cell Signaling #9204S |
| 14-3-3 pan | Immunoblot, immunoprecipitation | Santa Cruz #sc-1657 |
| Raptor | Immunoblot | Proteintech #20984-I-AP |
| Raptor | Immunoprecipitation | Thermo Fisher #42-4000 |
| Acetyl Lysine | Immunoblot, immunoprecipitation | Abcam #ab80178 |
| H3K9ac | Chromatin immunoprecipitation | Abcam #ab4441 |
| H3K27ac | Chromatin immunoprecipitation | Abcam #ab4729 |
| Cy3TM goat anti-rabbit IgG | Immunofluorescence | Invitrogen #A10520 |
| Cy3TM goat anti-mouse IgG | Immunofluorescence | Invitrogen #A10521 |
| Dnk pAB to goat IgG Alexa Fluor 488 | Immunofluorescence | Abcam #ab150129 |
| Goat pAb to Ms IgG (HRP) | Immunoblot | Abcam #ab97240 |
| Goat pAb to Rb IgG (HRP) | Immunoblot | Abcam #ab6721 |
| Rabbit IgG Isotype control | Immunofluorescence, immunoprecipitation | Abcam #ab172730 |
| Mouse IgG Isotype control | Immunofluorescence | Abcam #ab37355 |
